# Loss of Rab8a in B cells leads to increased antibody responses and class-switch recombination

**DOI:** 10.1101/2022.09.19.508414

**Authors:** Sara Hernández-Pérez, Alexey V. Sarapulov, M. Özge Balci, Eleanor Coffey, Akihiro Harada, Pieta K. Mattila

## Abstract

Rab8a is a small GTPase with a wide range of reported functions in different cell types, including vesicle recycling, vesicle traffic to cilia, cell ruffling, migration, neurite outgrowth, Toll-like receptor signalling and T cell receptor docking at the immune synapse. However, the role of Rab8a in B lymphocytes has not been described to date. Here, using a conditional B cell-specific Rab8a knockout mouse model, we investigate the role of Rab8a both *in vivo* and *in vitro*. Rab8a KO mice present enhanced antibody responses to both T-dependent and T-independent immunisations. Rab8a KO cells showed normal BCR trafficking and antigen processing and presentation but however, increased class-switch recombination. While the early BCR signalling responses, such as proximal kinase activation and calcium-flux, were normal, the signalling via AKT and ERK1/2 was decreased. We propose that the lack of Rab8a alters cellular signalling leading to enhanced antibody responses and increased class-switch recombination potentially via downmodulation of the PI3K/AKT/mTOR pathway.

## Introduction

B lymphocytes (B cells) are an essential part of the adaptive immune system, as they play a vital role in the humoral immune responses. To produce antibodies, B cells need to recognise a specific antigen via their B cell receptor (BCR). Although the processes leading to B cell activation and antibody production are relatively well characterised, many cellular players remain unknown. Tight spatiotemporal control of the vesicular traffic inside and outside the cell is critical for maintaining cellular homeostasis and regulating cellular responses to different cues. The Rab GTPase family regulates the secretory and endocytic pathways and, by targeting membrane compartments, they define the structural and functional identity of intracellular organelles (*Stenmark*, 2009; *Stenmark & Olkkonen, 2001*; *Zerial & McBride*, 2001). More than 60 different Rab proteins have been found in humans, controlling a plethora of cellular processes. Nevertheless, the role of the small Rab GTPases in B lymphocytes, especially in regard to antibody production, remains poorly investigated.

Two Rab8 isoforms, Rab8a and Rab8b, encoded by different genes, have been identified in mammals. Rab8a is ubiquitously expressed whereas Rab8b is predominantly found in brain, spleen and testes (*Armstrong et al*., 1996). Rab8 has been linked to a plethora of functions in multiple cellular pathways and in coordination with different effectors (*Hattula et al*., 2006; *Peränen*, 2011; *Roland et al*., 2007). Rab8 has been reported to localise to the endosomal recycling compartments together with Rab11 and Arf6 to promote the polarised transport of newly synthesised proteins and mediate protrusion formation and cell shape remodelling (*Ang et al*., 2003; *Hattula et al*., 2006; *Peränen & Furuhjelm*, 2001; *Vidal-Quadras et al*., 2017). Rab8a maintains the cell polarity of intestinal epithelial cells *in vivo*, regulating the localisation of apical proteins, and Rab8a knockout (KO) mice present a severe phenotype caused by microvillus atrophy (*Sato et al*., 2007, 2014). Rab8 is also found in the ciliary membrane and plays a role in ciliogenesis together with Rab11 (*Knödler et al*., 2010; *Lu & Westlake*, 2021; *Westlake et al*., 2011). In addition, Rab8 promotes the turnover of focal adhesions and controls the directionality of cell migration and invasion (*Bravo-Cordero et al*., 2007, 2016)

However, there are only a handful of studies describing the role of Rab8 in immune cells, and reports are exceedingly scarce in B lymphocytes. In the Jurkat T cells, Rab8 has been shown to colocalise with IFT20 and Rab11 and, assessed by the expression of mutant forms of Rab8, it was implicated in TCR recycling and final docking to the immune synapse via its interaction with VAMP3 (*Finetti et al*., 2015). It has also been shown in T cells that Rab8 participates in the surface expression of CTLA-4 through LAX-TRIM binding (*Banton et al*., 2014). In macrophages, Rab8a interacts with PI3K in membrane ruffles and early macropinosome membranes, and regulates TLR signalling, inflammation and polarisation via the AKT/mTOR pathway (*Luo et al*., 2014, 2018; *Tong et al*., 2021; *Wall et al*., 2017, 2019). In B cells, Porter et al. showed that Rab8a mRNA stability increases upon CpG stimulation via PTB (polypyrimidine tract-binding protein) (*Porter et al*., 2008), suggesting that Rab8a might be required during immune activation-induced cellular processes, but its functional role remains unknown.

In this work, we comprehensively explored the role of Rab8a in B lymphocytes for the first time both *in vivo* and *in vitro*. We detected a strong colocalisation of Rab8a with the internalised antigen along the antigen processing route, suggesting a potential role of Rab8a in antigen trafficking or processing. Thus, we took advantage of a B cell-specific, conditional Rab8a KO model to investigate the impact of Rab8a in B cell development, B cell activation, and antibody responses. Although no defects were found in BCR trafficking or antigen presentation, interestingly Rab8a KO mice showed increased antibody responses to both T-dependent and T-independent antigens *in vivo* and increased class-switch recombination (CSR) *in vitro*. These data were further supported by increased activation-induced deaminase (AID), as well as IgG2b and IgG2c expression in an RNAseq dataset. We propose that the lack of Rab8a alters cellular signalling, decreasing the activation of PI3K/AKT/mTOR pathway and thus modulating AID expression and increasing CSR.

## Results

### Rab8a colocalises with the internalised antigen vesicles on the processing route

In our previous studies, we reported the heterogenous colocalisation of several endosomal Rab proteins (Rab5, Rab6, Rab7, Rab9 and Rab11) with the internalised antigen (*Hernández-Pérez et al., 2020*). To gain more understanding into the workshare of different Rab proteins, we investigated the Rab protein expression profiles (*ImmGen Datasets*) and identified Rab8a with relatively high expression level in B cells. A previous report in T cells showing that Rab8 participates in TCR transport (*Finetti et al*., 2015), prompted us to study the colocalisation of the internalised antigen-BCR complexes with Rab8a. A20 D1.3 cells and primary B cells were activated with fluorescent anti-IgM as a surrogate antigen, fixed at different time points, and stained for Rab8a. We detected a clear and fairly constant colocalisation of Rab8a with the internalised antigen (Fig. 1A and Fig. S1A). Shortly after internalisation, Rab8a colocalised with the antigen close to the plasma membrane, while at later time points, Rab8a trafficked together with the antigen to the perinuclear compartment (Fig. 1A). A similar phenomenon was also observed in primary B cells isolated from the spleen of C57BL/6N mice (Fig. 1A). These data suggested that Rab8a might play a functional role in the antigen processing either at the stage of BCR-antigen internalisation or antigen processing and presentation.

**Figure 1:**
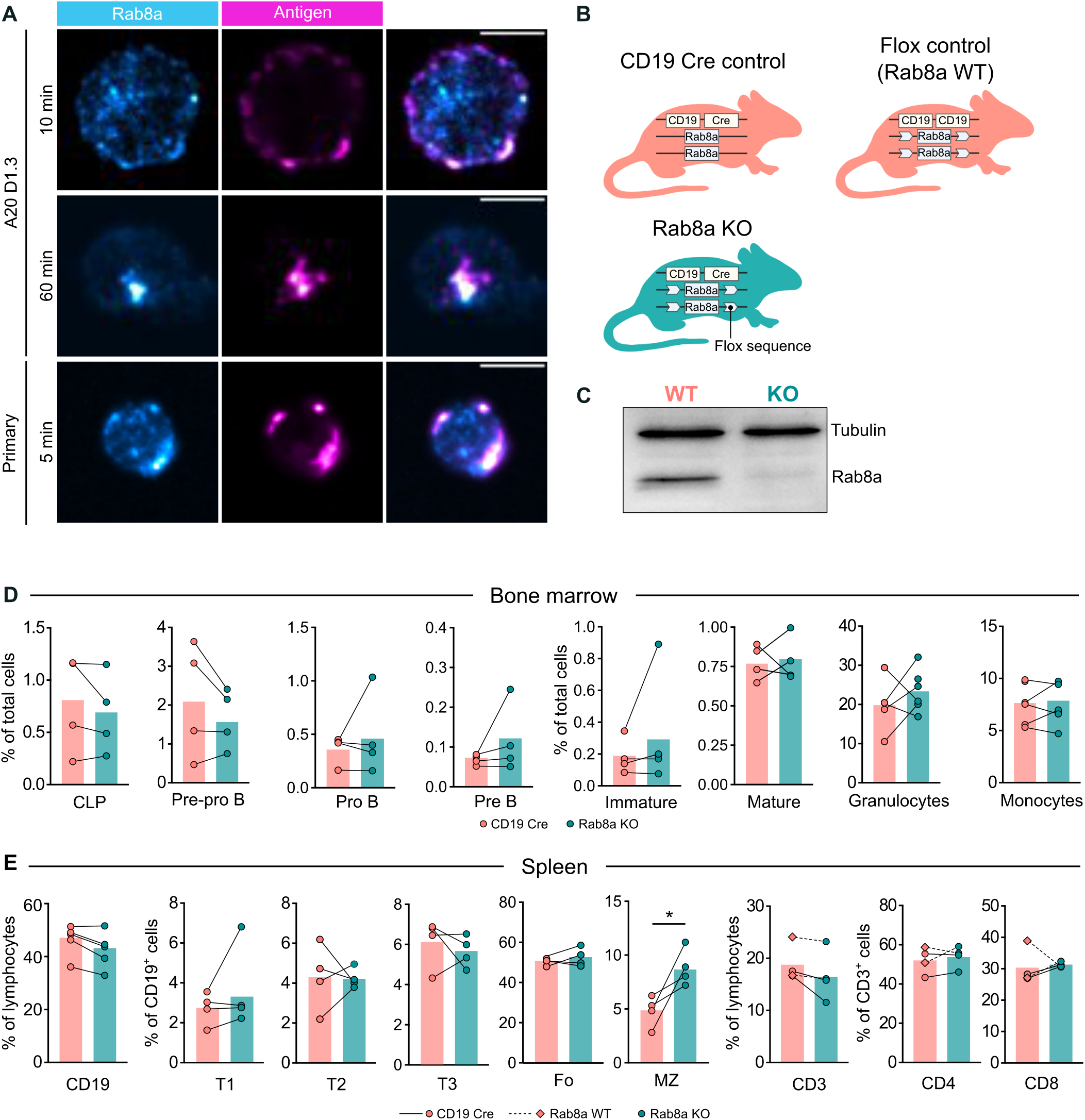
Establishment of a conditional B cell Rab8a KO mice and immunophenotyping. **(A)** A20 D1.3 (top panels) or primary B cells (bottom panel) were activated with soluble surrogate antigen (anti-IgM, Magenta Hot LUT) and stained with anti-Rab8a (Cyan Hot LUT). Merged images are shown on the right. Images were acquired with a spinning disk confocal microscope (SDCM). One plane (2D) is shown. Scale bar: 5 μm. **(B)** Different mouse models used in the present study. The boxes represent the CD19, Cre and Rab8a genes. The arrows represent the flox sequences. **(C)** WB showing Rab8a expression in primary B cells isolated from the spleens of Rab8a WT and KO mice. See also Fig. S1B. **(D)** Bone marrow and **(E)** spleen cell populations from WT or CD19 Cre controls and Rab8a KO mice were extracted for analysis by flow cytometry (immunophenotyping). Data are shown as mean values (bar) of 4-5 independent experiments (every dot represents one mouse). Mice pairs (same experimental date) are connected by a line. Statistics: paired t-test.

### B cell development is largely normal in Rab8a^-/-^ mice

To study the role of Rab8a in the B cell immune responses *in vivo*, we generated a B cell conditional Rab8a^-/-^ knock out (KO) by crossing the Rab8a^flox/flox^ strain (Sato et al., 2007) with CD19-Cre knock-in mice (*Rickert et al*., 1997) (Fig. 1B). In the heterozygous CD19-Cre mice, the endogenous CD19 gene (starting in pro-B stage) is disrupted by the Cre gene, leading to a 90-95% Cre-mediated deletion efficiency of the floxed gene in splenic B cells (*Rickert et al*., 1997). Successful establishment of the model was confirmed by PCR (*data not shown*) and western blot (Fig. 1C and Fig. S1B). B cells isolated from the spleen of Rab8a KO mice showed minimal Rab8a expression compared to their WT counterparts. In addition, Rab8b expression remained unmodified, confirming the specificity of the deletion and demonstrating that Rab8b expression levels do not increase to compensate for the loss of Rab8a (Fig. S1B).

To evaluate the potential role of Rab8a in BCR trafficking, we evaluated the surface expression and internalisation of the IgM BCR in WT and Rab8a KO cells. Rab8a KO cells showed no difference in the levels of surface IgM BCR nor the levels of surface MHCII, required for antigen presentation (Fig. S1C). We also investigated the internalisation of the BCR both in steady-state and after BCR-mediated activation, but no differences were detected (Fig. S1D, E). To evaluate the effect of the Rab8a deletion on lymphocyte development, we extracted cells from bone marrow (Fig. 2C), spleen (Fig. 2D) and lymph nodes (Fig. S2A) and analysed the different immune cell populations as previously described (*Sarapulov et al*., 2020). To fully match the gating of the B cell (CD19+) populations, CD19-Cre controls were used here, instead of the littermate WT flox controls (see Fig. 2A) so that all the cells had the same level of CD19 (*Yasuda et al*., 2021). We found no significant differences between WT and Rab8a KO mice, except for a slight increase in the frequency of marginal zone (MZ) B cells (Fig. 2D), suggesting that the lack of Rab8a does not compromise B cell development.

**Figure 2:**
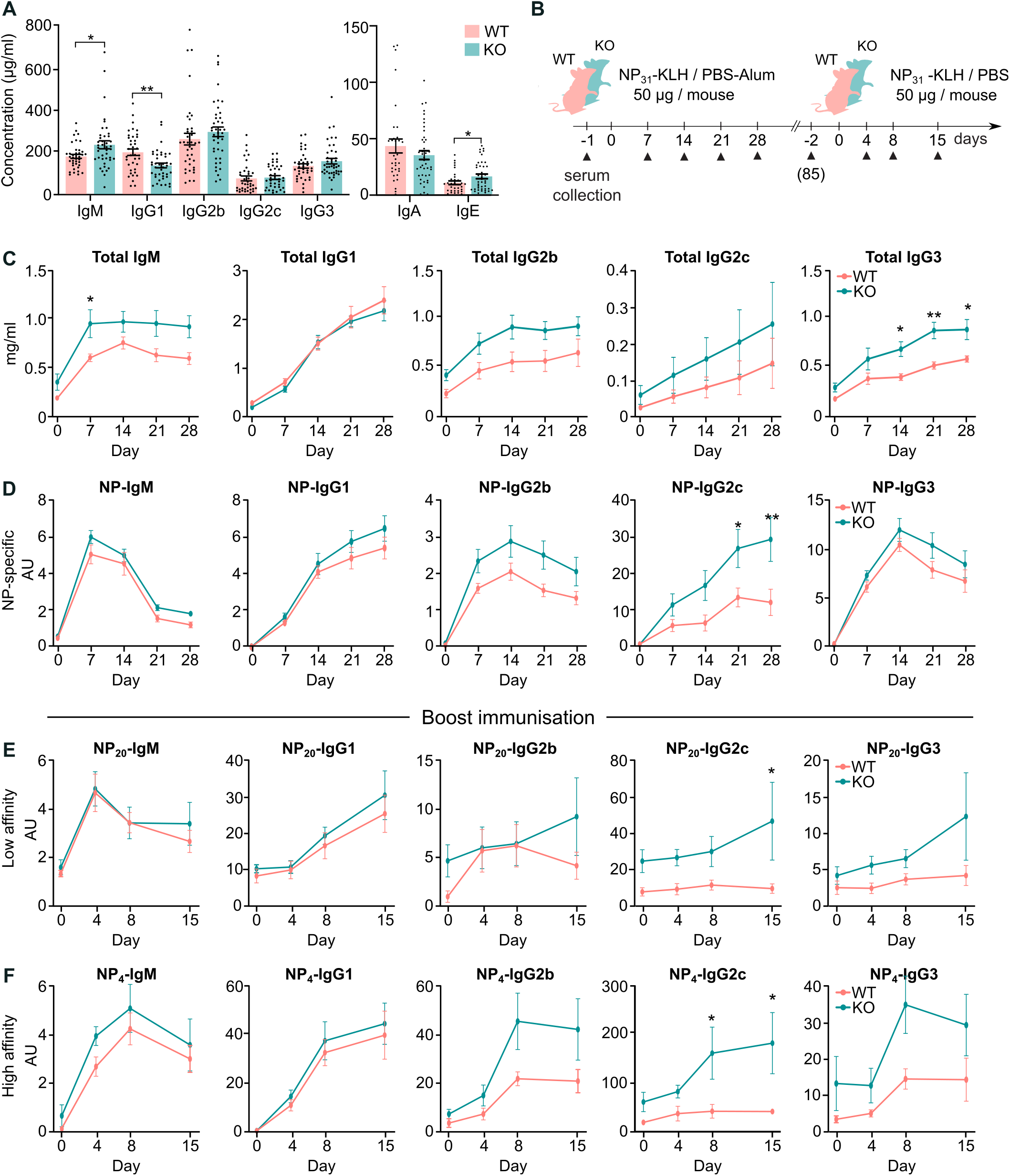
Altered basal antibody levels and increased T-dependent immune responses Rab8a KO mice. **(A)** Basal antibody levels obtained from the sera of non-stimulated mice (n = 35-45 mice). Statistics: unpaired t-test. **(B)** Schematics for the NP-KLH immunisation schedule. **(C)** Total levels of antibodies after primary NP-KLH immunisation. **(D)** NP-specific levels (NP20) after primary NP-KLH immunisation. **(E)** Low-affinity NP-specific (NP20) antibody levels after boost immunisation. **(F)** High-affinity NP-specific (NP4) antibody levels after boost immunisation. **(C-F)** n = 5 mice per group. Mean ± SEM. Statistics: 2-way ANOVA (Sidak’s multiple comparisons test).

### Rab8a KO mice show increased antibody responses *in vivo*

Next, we evaluated the *in vivo* immune responses in these mice. First, blood was collected from the mice, and the basal antibody levels in the serum were analysed using ELISA. We detected slightly increased IgM and IgE levels and decreased IgG1 levels in the serum of the Rab8a KO mice, while all other isotypes remained unchanged (Fig. 2A).

To investigate the response to T-dependent antigens, mice were immunised with the hapten NP conjugated to KLH in the presence of Imject^™^ alum (NP-KLH; experimental procedure in Fig. 2B). Unexpectedly, Rab8a KO mice showed an overall increased response to immunisation (Fig. 2C, D), especially toward IgG2c and IgG3 isotypes. After the boost immunisation, Rab8a KO showed increased responses in both high- (Fig. 2E) and low- (Fig. 2F) affinity antibody responses, especially toward IgG2c and IgG3. We further dissected the antibody response in these mice by immunising them with NP-KLH and looking at the germinal centre responses. Germinal centres are highly dynamic structures: they are known to peak around 5 to 10 days after immunisation, decreasing after that (*Victora & Nussenzweig*, 2012). Hence, we immunised the mice and collected the spleens on day 9 (expansion phase) and day 21 (contraction phase) after immunisation to analyse the kinetics of the response. We detected a robust induction of GC B cells (CD95^+^ GL-7^+^) following the immunisation (day 9), but no differences were observed between the WT and Rab8 KO mice (Fig. S2B, C). Consistent with the kinetics of the GC reaction, we detected a decrease of around 50% in the percentage of GC cells after 3 weeks (day 21), with no differences between WT and KO (Fig. S2B). The isolated splenocytes were also plated on NP-coated ELISpot plates to detect the number of NP-specific antibody-secreting cells (ASCs), but no differences were detected at day 9 (Fig. S2D).

Next, we studied the antibody responses after immunisation with a T-independent type II (NP-Ficoll; Fig. 3A-C) or type I (NP-LPS; Fig. 3D-E) antigens. Similarly to the T-dependent antigen, Rab8a KO mice produced more IgG antibodies, especially of the IgG1, IgG2c and IgG3 subclasses (Fig. 3). We also followed the numbers of ASCs using ELISpot 9 days after NP-FICOLL immunisation, but no differences were detected between WT and KO mice (Fig. S2E).

**Figure 3:**
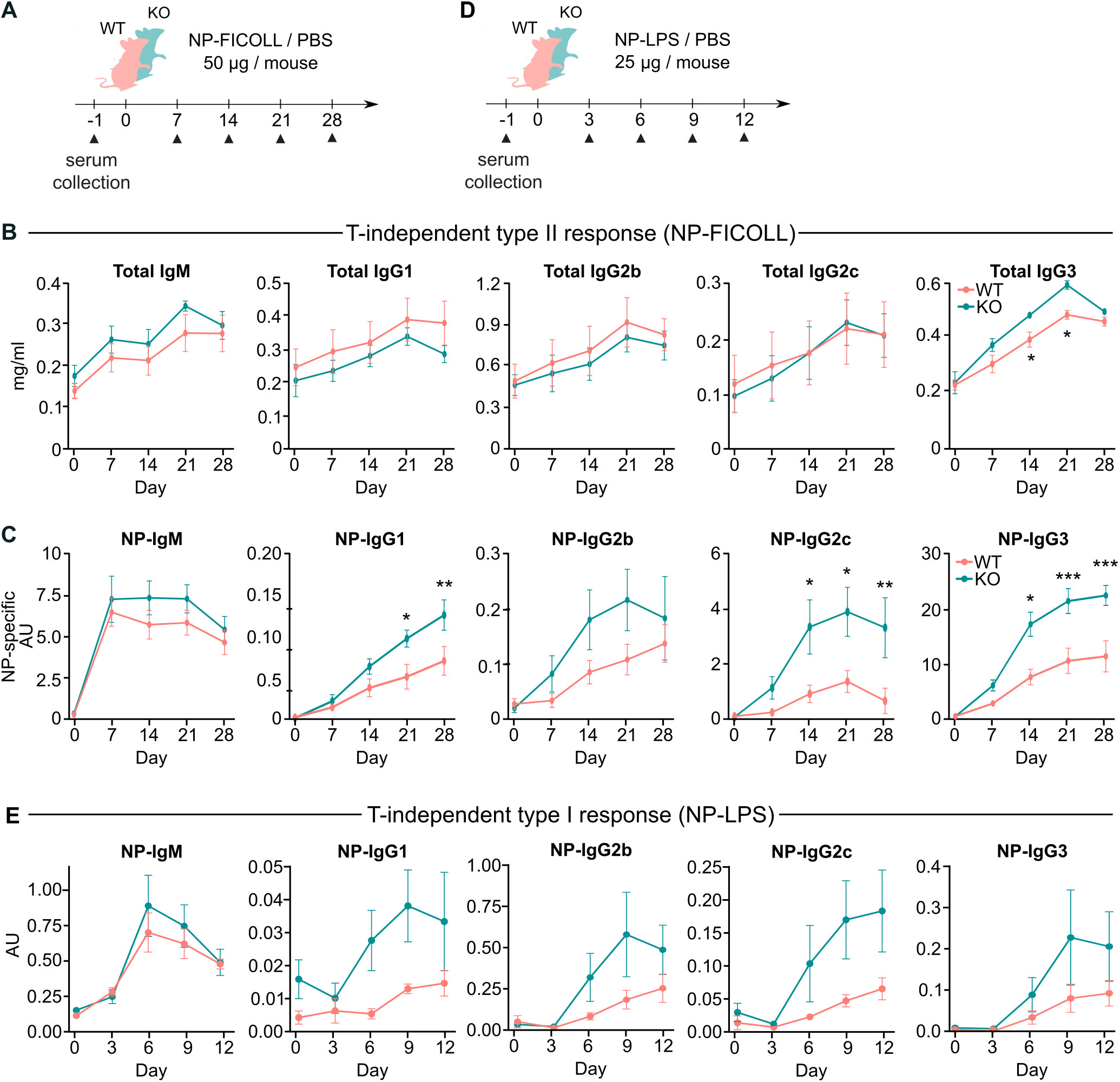
Increased T-independent antibody responses in Rab8a KO mice. **(A)** Schematic representation of the NP-FICOLL immunisation schedule. **(B)** Total levels of antibodies and **(C)** NP-specific antibody levels (NP20) after NP-FICOLL immunisation. **(D)** Schematic representation of the NP-LPS immunisation schedule. **(E)** Total levels of NP-specific antibodies (NP20) after NP-LPS immunisation. (B, C, E) n = 5 mice per group. Mean ± SEM. Statistics: 2-way ANOVA (Sidak’s multiple comparisons test).

### Rab8a KO cells upregulate AID and class-switched immunoglobulin isotypes upon activation

To gain more information about the possible mechanism mediating the increased antibody responses detected *in vivo*, we performed RNA sequencing (RNAseq) in resting or *in vitro* activated B cells. B lymphocytes were isolated from the spleen or lymph nodes and either directly lysed for RNA isolation (resting samples, spleen or lymph node) or activated the spleen-derived B cells with anti-IgM/OVA beads and co-cultured with OT-II cells, to allow for T cell help, for 72 h prior to B cell sorting (activated samples). The gene profile of each sample was depicted using a principal component analysis (PCA; Fig. S3A) and the Pearson correlation between each sample (Fig. S3B). In the PCA, the largest variability between different samples originated from the tissue (spleen/lymph node) and the activation state (resting/activated), rather than from Rab8a expression (WT/KO).

Next, we compared the differentially expressed genes (DEG) in the WT and KO samples in different conditions (Fig. 4A-B and Fig. S3C-D). We found few DEGs in the Rab8a KO mice compared to the WT mice (17 in spleen, 10 in lymph nodes and 15 in activated samples). Of these, only 3 were consistent through all conditions: GPRC5b was upregulated, and the pseudogenes Gm4864 and Gm10036 were downregulated in Rab8a KO B cells. Interestingly, among the 15 DEGs between the activated WT and KO cells, we found a significant increase in AID (Aicda), IgG2b (Ighg2b) and IgG2c (Ighg2c), in line with the elevated IgG2c response to T cell-dependent immunisation observed *in vivo* (Fig. 4B). In addition, we also observed a decrease in CD138 (Sdc1) (Fig. 4B). Next, the 15 DEGs found in the activated Rab8a KO/WT cells were analysed using Gene Ontology (GO) analysis (Fig. 4C). GO results showed that DEGs in the KO mice were enriched in the following functional categories: inflammatory response, production of molecular mediators, immunoglobulin production and cytokine production. Consistent with the *in vivo* findings, these results suggest that Rab8a KO mice are prone to mount increased immune responses as reflected by the increased antibody titers following the immunisations (see Fig. 2 and 3).

**Figure 4:**
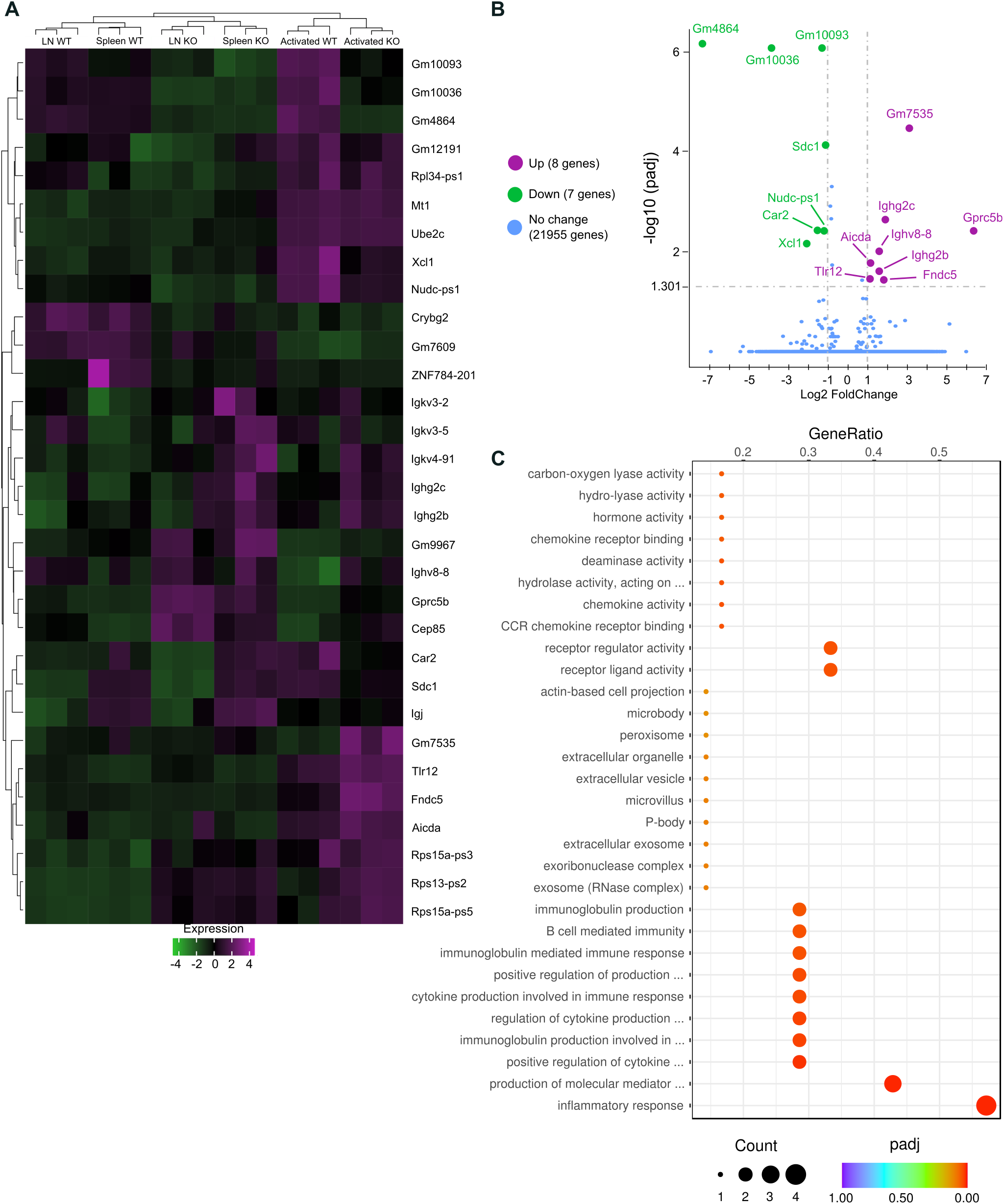
Rab8a KO cells show an increase activation profile. **(A)** Heat map showing the DEGs in Rab8a WT and KO samples from spleen (activated and non-activated) and lymph nodes (non-activated). Purple colour shows upregulation and green colour shows downregulation in Rab8a KO B cells compared to WT cells. **(B)** Volcano plot showing the 15 DEGs found in activated samples (splenic B cells). Purple colour shows upregulation and green colour shows downregulation in Rab8a KO B cells compared to WT cells. **(C)** GO analysis of the 15 DEGs found in the activated samples.

### Rab8a KO cells show increased class-switch recombination *in vitro*

To better understand the altered immune responses in Rab8a KO mice, we next evaluated B cell proliferation, immunoglobulin class-switching and antibody secretion in cells activated *in vitro*. First, we checked the proliferation of B cells in response to different stimuli. B cells were isolated from the splenocytes and activated *in vitro* for 72 hours using anti-IgM, LPS, CD40L, CpG or a combination of those in the presence of IL-4. We found no differences in B cell proliferation in response to any of these stimulations (Fig. 5A). However, the Rab8a KO cells showed increased percentage of class-switched IgG1 and IgG2c cells, and decreased numbers of plasma cells (CD138^+^ cells) (Fig. 5B). The reduction in CD138^+^cells was in agreement with the RNA sequencing data showing decreased expression of CD138 (Sdc1) in Rab8a KO cells (see Fig. 4B). Despite the decreased numbers of CD138^+^ cells, the amount of secreted IgG1 was higher in the culture supernatants of Rab8a KO cells stimulated with combination of F(ab’)_2_ anti-IgM, LPS, IL-4 and IL-2, as compared to the WT counterparts, but no difference was observed with LPS + IL-4 alone. The data suggested that BCR-mediated signalling may profoundly influence response outcome imposed by other activatory signals in the Rab8a KO cells (Fig. 5C).

**Figure 5:**
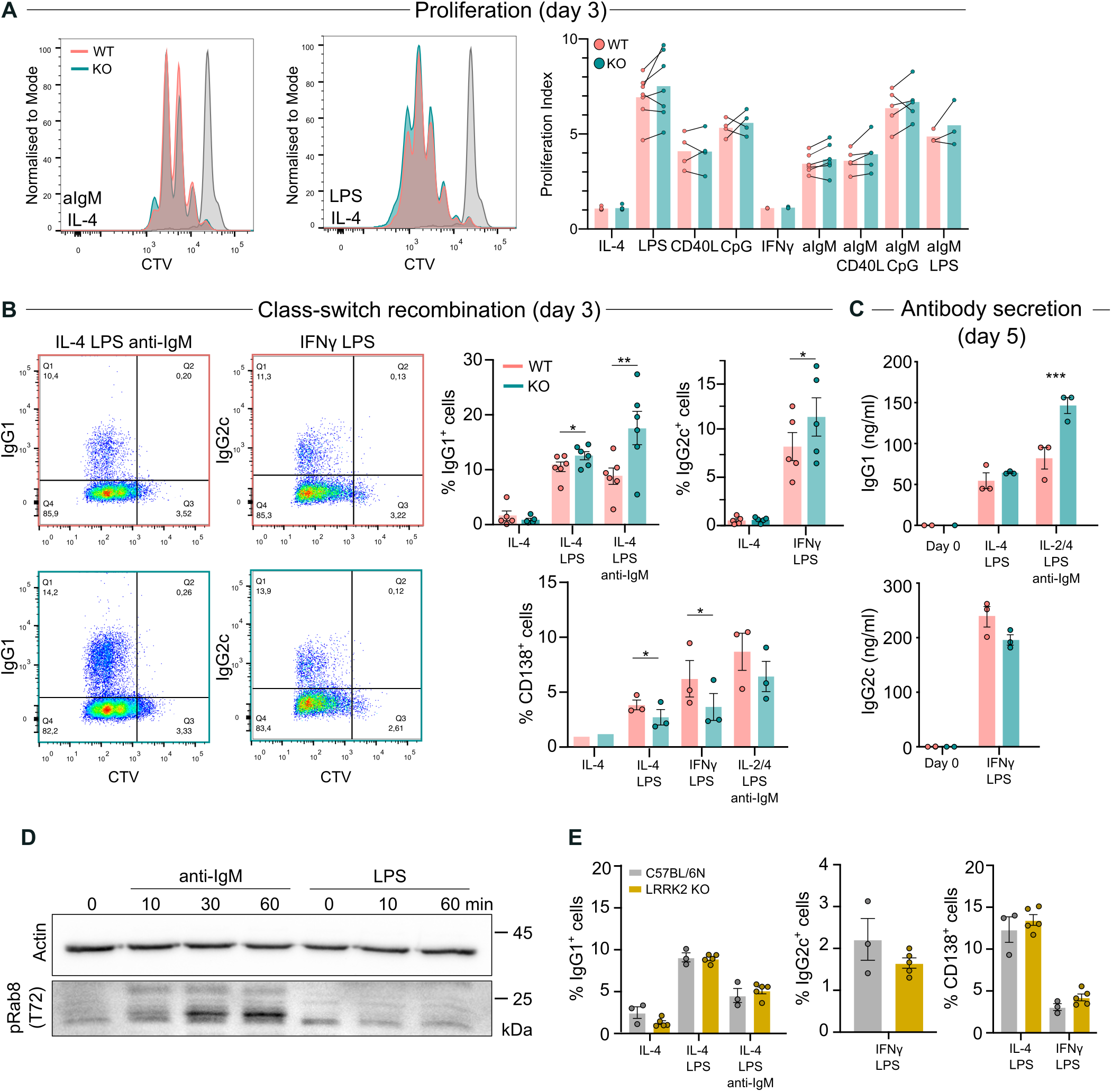
Rab8a KO mice exhibit increased class-switch recombination in *vitro*. **(A)** Analysis of WT or Rab8a KO B cell proliferation in response to different stimuli. Mouse primary B cells were loaded with CTV, and proliferation was analysed after 72 h with flow cytometry. Representative plots of F(ab’)_2_ anti-IgM and LPS are shown. Proliferation responses (mean bar) are measured as proliferation index (n = 3-7 independent experiments). Statistics: paired t-test, no differences. **(B)** Analysis of WT or Rab8a KO B cell class-switch (IgG1^+^, IgG2c^+^) and plasma cell differentiation (CD138^+^) in response to different stimuli after day 3. Data are shown as representative flow cytometry plots (left) and percentage of cells (mean ± SEM; right) from n = 3-6 independent experiments. **(C)** Analysis of antibody secretion, measured by ELISA, in WT and Rab8a KO cells stimulated as in (B) for 5 days. Data are presented as mean ± SEM of n = 3 independent experiments. **(D)** Western blot showing Rab8a phosphorylation (T72) after BCR (10 μg/ml F(ab’)_2_ anti-IgM) or TLR (10 μg/ml LPS) stimulation in A20 D1.3 cells. Representative blot; quantification is shown in Fig. S4A. **(E)** Analysis of WT or LRRK2 KO mouse primary B cell class-switch and plasma cell differentiation in response to different stimuli after day 3. Data are presented as mean ± SEM of n = 3 independent experiments.

The distinct altered responses of Rab8a KO cells to stimulations with BCR agonists, prompted us to address whether Rab8 could get activated upon BCR stimulation. In addition to the regulation by the GTP/GDP status, the activity of Rab proteins can also be modulated by phosphorylation, and, for instance, it has been shown that Rab7 gets phosphorylated upon BCR signalling (*Satpathy et al*., 2015) Thus, we examined the potential phosphorylation of Rab8 upon BCR stimulation and, additionally, upon TLR stimulation as it has been reported that Rab8a is phosphorylated upon TLR activation in macrophages (*Luo et al*., 2014). Notably, we detected robust phosphorylation of Rab8a on the threonine 72 (pT72) following BCR (F(ab’)_2_ anti-IgM) stimulation in A20D1.3 mouse B cells (Fig. 5D and Fig. S4A). However, no response to TLR (LPS) was detected. We further verified that Rab8 phosphorylation also occurred in A20 D1.3 cells upon stimulation with hen egg lysozyme (HEL), the monovalent antigen recognised by the D1.3 IgM BCR (Fig. S4B). In addition, since the anti-pT72 antibody is known to cross-react with other Rab proteins, such as Rab10 (*Lis et al*., 2018), we verified the specificity of Rab8a phosphorylation by overexpressing a citrine-Rab8a fusion protein in A20 D1.3 cells. We found an increase also in the phosphorylation of citrine-Rab8 verifying the specificity of the phosphorylation, although to a lesser extent than seen for the endogenous protein (Fig. S4C).

To address the role of Rab8 phosphorylation on the anti-body responses, we targeted the protein leucine-rich repeat kinase 2 (LRRK2). It has been established that Rab8, as well as other Rab proteins, can be phosphorylated by LRKK2, although the role of phosphorylation remains unclear (*Eguchi et al*., 2018; *Madero-Pérez et al*., 2018; *Steger et al., 2017*). Thus, we analysed the levels of Rab8a phos-phorylation in WT primary B cells treated with two different LRRK2 inhibitors, GSK2578215A (GSK) and GNE-7915 (GNE) (*data not shown*) and using primary B cells isolated from LRRK2 KO mice (Fig. S4D). Notably, we observed that LRRK2 inhibition or knock-out decreased but did not completely abolish Rab8 phosphorylation. Next, we addressed the possible effects of diminished Rab8 phosphorylation in CSR assay using the LRRK2-inhibited (Fig. S5A-C) and the LRRK2 KO primary cells (Fig. 5E). However, we found no differences in the percentage of class-switched cells and plasma cells. We also quantified the basal antibody titers in the serum of LRRK2 KO mice, but Statistics: paired t-test.they did not display the same phenotype as Rab8 KO mice (Fig. S5D, compared to Fig. 2A). All together, these results suggest that while Rab8a modulates CSR, this is independent on the phosphorylation by LRRK2.

### Rab8a KO mice present unaltered antigen presentation and migration

Inspired by the published reports in epithelial cells, where Rab8 plays a role in cell migration (*Bravo-Cordero et al*., 2016), next, we investigated B cell migration *in vitro* and *in vivo*. However, no differences were found in migration towards CXCL12 nor CXCL13 using a transwell assay (Fig. 6A), nor in the *in vivo* migration to the lymph nodes (Fig. S6A). We next evaluated the antigen presentation potential of the Rab8a KO cells using the *in vitro* Eα peptide presentation system but found normal levels of peptide presentation (Fig. 6B). As the Eα system only evaluates the peptide-MHCII loading capacity of the cells, but not the antigen processing, we also studied the antigen presentation by taking advantage of the OT-II system. B cells from Rab8a KO mice or the littermate controls were purified and stimulated *in vitro* with anti-IgM/ovalbumin (OVA) beads and co-cultured with T cells isolated from OT-II mice able to recognise OVA peptides. After 72 h cell proliferation and cytokine secretion were analysed. IL-2 is secreted by the activated T cells, while activated B cells produce IL-6, a proinflammatory cytokine that promotes T follicular helper (TFH) cell differentiation. We did not observe significant changes in T or B cell proliferation, nor in IL-2 or IL-6 secretion, although they all showed a minor trend for upregulated responses with Rab8a KO cells (Fig. 6 C-F). Altogether, these data suggest that Rab8a is not required for normal antigen presentation, B cell proliferation or migration.

**Figure 6:**
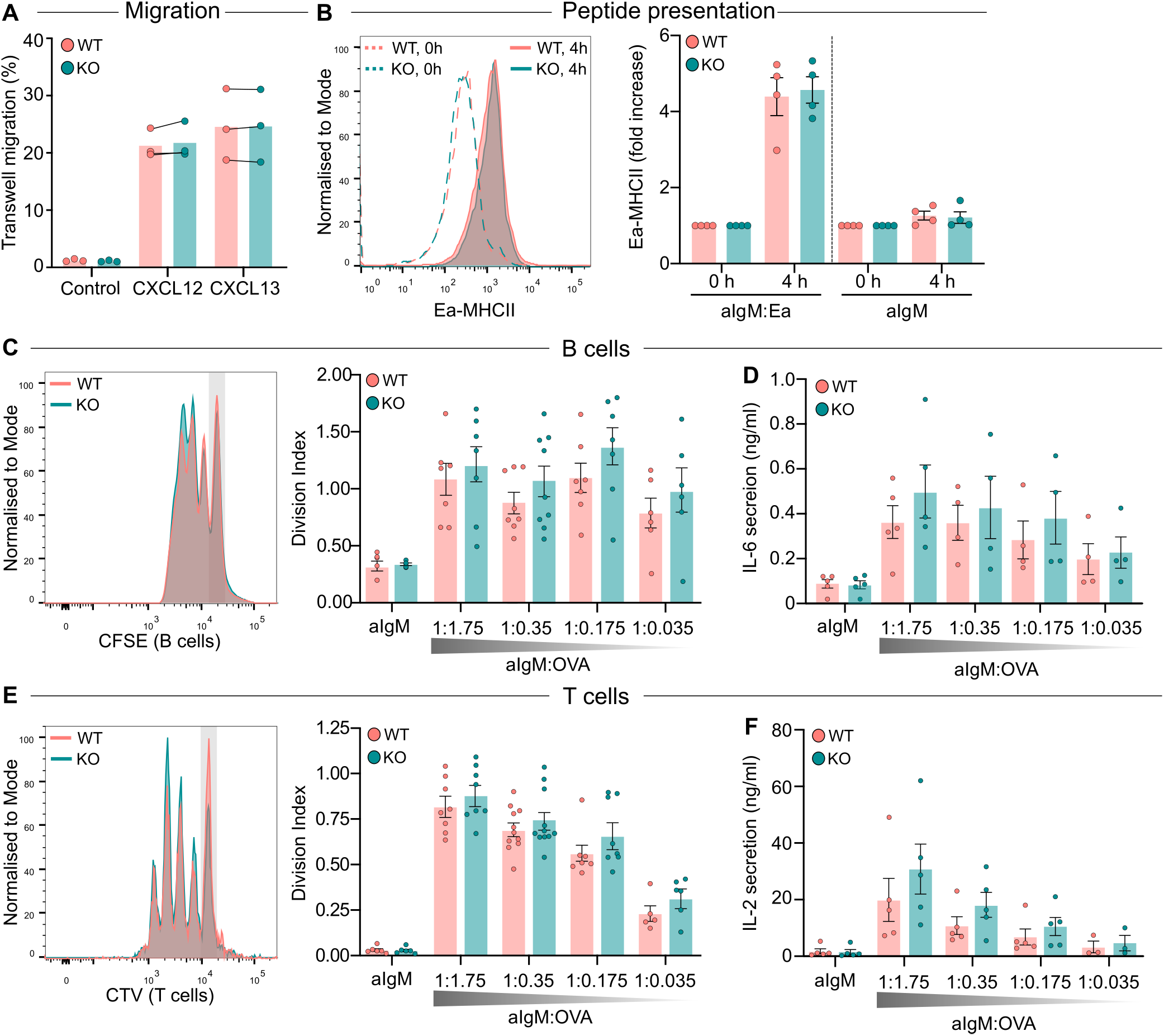
Rab8a KO mouse shows normal migration, antigen presentation and B and T proliferation. **(A)** Transwell migration after 4 h in response to CXCL12 or CXCL13 using primary WT and Rab8a KO cells isolated from the spleen. Paired t-test, no differences (n = 3 independent experiments; each dot one mouse). **(B)** Primary WT and Rab8a KO B cells were activated *in vitro* with anti-IgM/Ea peptide-coated beads (aIgM:Ea) and peptide presentation was measured using flow cytometry after 4 hours. As a negative control, beads coated with anti-IgM alone were used (aIgM). Representative plots (mean ± SEM) and data from 4 individual experiments are shown as fold increase of the MHCII-Eα signal after 4 hours, compared to 0 hours (arbitrary value of 1). **(C-F)** Primary WT and Rab8a KO B cells were labelled with CFSE and OT-II T cells were labelled with CTV. B cells were activated *in vitro* with anti-IgM/OVA-coated beads (aIgM:OVA, in different ratios) and co-cultured with the labelled OT-II T cells for 3 days. As a negative control, beads coated with anti-IgM alone were used (aIgM). After 3 days, proliferation was analysed using flow cytometry. Representative plots and division index of **(C)** CFSE-labelled B cells and **(E)** CTV-labelled T cells (mean ± SEM; n > 6 independent experiments) are shown. **(D)** Amount of IL-6 and (F) IL-2 in the T-B cell co-culture supernatants after 3 days measured by ELISA (mean ± SEM; n = 4-5 independent experiments). Paired t-test.

### Rab8a KO B cells show normal proximal tyrosine kinase signalling, but altered downstream signalling in response to BCR stimulation

Finally, we investigated whether Rab8a could modulate BCR signalling. We studied early BCR signalling in response to soluble and surface-bound anti-IgM. Rab8a KO B cells seeded on anti-IgM coated glass formed symmetrical immunological synapses (Aspect Ratio), and no differences in spreading (area at the contact site) or signalling (pPLCγ2 / pBtk) were detected (Fig. 7A and Fig. S7A). Next, we went on to study proximal tyrosine kinase signalling (pSyk / pLyn) in response to soluble (Fig. 7B) and surface-bound (Fig. S7B) F(ab’)_2_ anti-IgM by Western blotting but saw no differences. We also analysed the calcium flux in response to soluble F(ab’)_2_ anti-IgM using flow cytometry and found Rab8a KO cells behaving similarly to the WT counterparts (Fig. 7C). Next, we investigated if the downstream BCR signalling pathway could be altered in Rab8a KO cells in response to BCR activation (Fig. 7D). We detected a slight decrease in both AKT and ERK signalling in Rab8a KO cells and an alteration in the kinetics of PI3K signalling. A slight but not significant decrease in mTOR signalling, assessed by phosphorylation of S6, was observed. Altogether, these data suggested that while loss of Rab8a does not compromise proximal BCR signalling it leads to modulated downstream signalling pathways in response to BCR stimulation.

**Figure 7:**
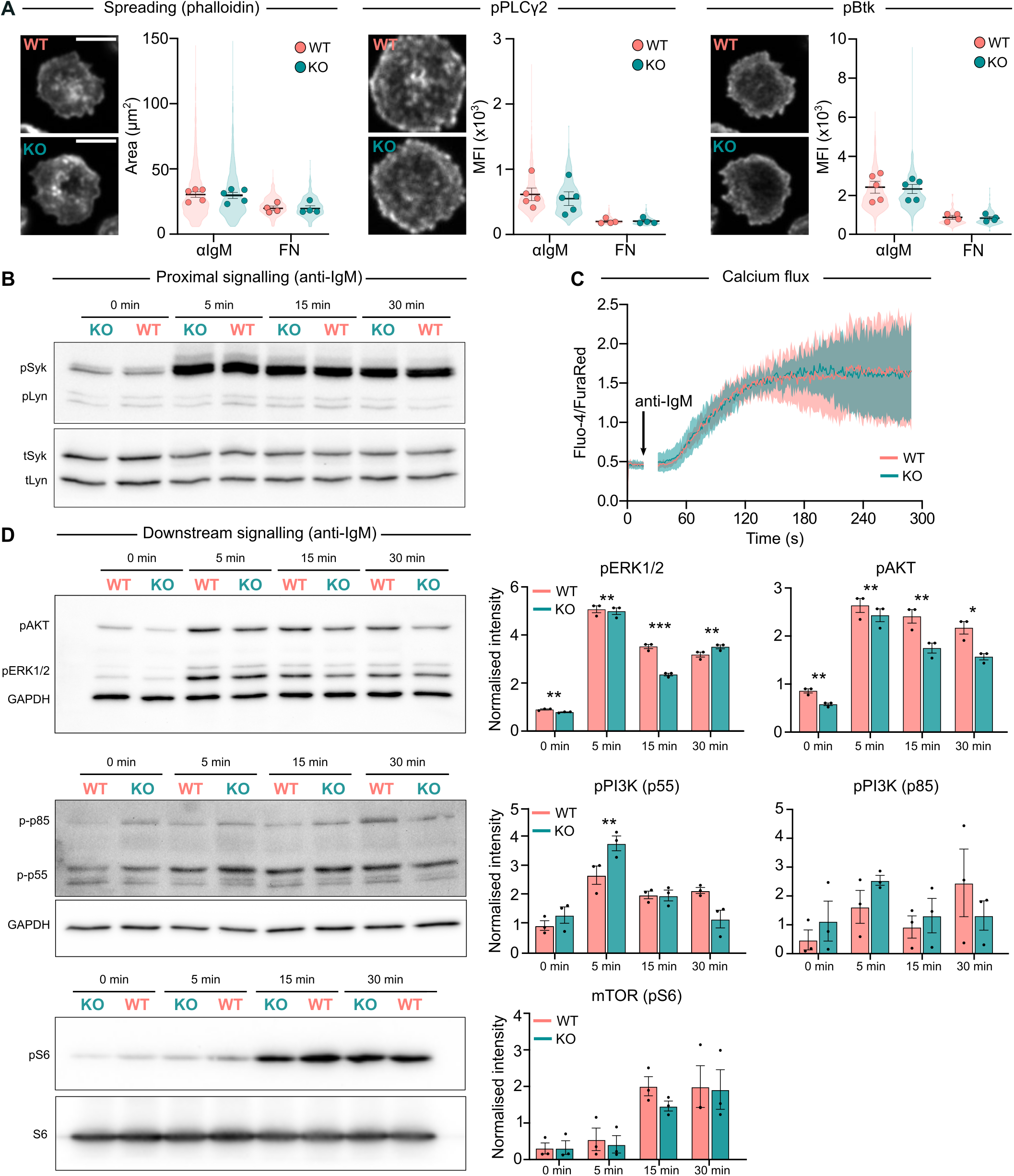
Altered downstream BCR signalling in Rab8a KO B cells. **(A-B)** Analysis of B cell activation in response to anti-IgM coated glass. Primary B cells from WT or Rab8a KO mice were seeded on anti-IgM coated glass for 15 minutes, fixed and stained with phalloidin (actin) and phospho-antibodies. Cells were imaged using a spinning disk confocal microscope. Spreading response was measured as the cell area detected on phalloidin channel and BCR signalling was measured by phospho-PLCγ2 and phospho-Btk (MFI). The violin plots represent the distribution of the population (all analysed cells, >100 cells per experiment) and dots represent the mean of individual experiments (n = 4-5). **(B)** Analysis of proximal BCR signalling by Western blot. Cells were activated with 10 μg/ml of F(ab’)_2_ anti-IgM in solution for 5, 15 and 30 minutes. Statistics: paired t-test. **(C)** Calcium flux was measured in primary B cells isolated from WT and Rab8a KO mice, loaded with Fluo-4 and FuraRed, in response to 5 μg/ml of F(ab’)_2_ anti-IgM. The response is measured as the ratio between Fluo-4 and FuraRed GeoMean at each timepoint. The thick line represents the mean, and the shadow area represents the SD of n = 4 independent experiments. The arrow marks the addition of anti-IgM. **(D)** Analysis of downstream BCR signalling in response to F(ab’)_2_ anti-IgM anti-IgM by Western blot. Cells were activated with 10 μg/ml of F(ab’)_2_ anti-IgM in solution for 5, 10 and 15 minutes, lysed and analysed for ERK1/2, AKT, PI3K and mTOR (S6) signalling. N = 3 independent experiments.

## Discussion

While all cells need a tight regulation of the endosomal traffic, this is imperative in immune cells performing specialised functions. B lymphocytes require an exquisite regulation of their different intracellular pathways as these are involved in, among other things, BCR-antigen internalisation, antigen processing and presentation, cytokine secretion and antibody production. Due to the prominent role of Rab GTPases in vesicle trafficking, it comes as no surprise that these proteins play several roles in regulating the immune responses (*Krzewski & Cullinane*, 2013). However, only a few reports have shown a direct link between Rab GTPases and antibody responses (*Lam et al*., 2016; *Pone et al*., 2015; *Satpathy et al*., 2015). In this study, we focused on the B cell immune responses and the role of Rab8a, a small GTPase with a wide range of functions reported in different cell types. To note, the role of Rab8a in B lymphocytes has not been described to date and our results position Rab8a as a regulator of B cell class-switching and antibody responses, presumably via a modulatory function in of the PI3K/AKT/mTOR pathway, rather than an effect on the BCR/antigen processing pathway.

We found Rab8a to colocalise with the internalised antigen vesicles along the antigen processing route in primary cells and in the A20 D1.3 cell line (Fig. 1A). To study the possible role of Rab8a in the antigen processing and the immune response, we generated a conditional KO mouse model lacking Rab8a in B cells (CD19^Cre/+^, Rab8a^flox/flox^). The deletion of Rab8a did not significantly alter B cell development, except for a slight increase in the numbers of MZ B cells (Fig. 1C-D). As Rab GTPases typically regulate vesicular compartments, we performed a set of *in vitro* experiments to test if Rab8a would regulate antigen internalisation, processing, or presentation. However, we found no significant alterations in Rab8a KO cells in assays for BCR internalisation, Eα peptide presentation or antigen (OVA) processing and presentation to cognate T cells, suggesting that Rab8a does not critically regulate the BCR/antigen trafficking routes (Fig. S1C, D and Fig. 6). Knock-down of Rab8a did not affect the expression level of Rab8b (Fig. S1B). As compensation of Rab8a function by other Rab family proteins, due to the specialized functions of Rab proteins, is in general unlikely, we did not investigate the expression levels of other Rab proteins in our mouse model. However, for instance, Rab11 is known to colocalise with Rab8 in recycling endosomes and both proteins act together to regulate the trafficking of these vesicles (Knödler et al., 2010). Thus, it would be interesting to investigate the possible redundancy between Rab8 and Rab11 or other Rab proteins on BCR/antigen trafficking in the future.

Interestingly, even though Rab8a KO did not alter BCR/antigen process or presentation, we observed altered antibody responses. When we analysed the antibody levels in the WT and Rab8a KO mice, we found increased basal levels of IgM and IgE, but decreased IgG1 in the sera of Rab8 KO mice (Fig. 2A). Interestingly, upon immunisation with TD and TI antigens, we also observed increased NP-specific antibody levels at different IgG isotypes (Fig. 2 and 3). Aiming to gain insights into the possible mechanisms behind the increased antibody responses, we performed bulk RNAseq of the WT and Rab8a KO B cells from lymph nodes and spleen in resting state or after activation *in vitro* (Fig. 4 and Fig. S3). We found only three DEGs consistently upregulated (GPRC5b) or downregulated (the pseudogenes Gm4864, Gm10036) throughout the different conditions. Of these, GPRC5b, an orphan G-protein-coupled receptor (GPCR) linked to obesity and inflammation, was deemed as an interesting hit (*Hirabayashi & Kim*, 2020; *Zambrano et al*., 2019). However, this was not further pursued as we could not validate the GPRC5b upregulation at the protein level using western blot (*data not shown*). Interestingly, when analysing the *in vitro* activated cells, increased expression of AID, IgG2b and IgG2c genes was observed, well in line with the increased class switching to the IgG2b, IgG2c, and IgG3 secretion in the immunised mice *in vivo* (Fig. 2 and 3). However, also decreased levels of syndecan-1 (CD138), a marker of plasma cells was observed (Fig. 4). We then analysed the percentage of class-switched cells upon *in vitro* stimulation of splenic B cells. Rab8a KO cells showed increased percentage of class-switched cells, especially in those conditions where the BCR was stimulated (Fig. 5A-C). However, in these *in vitro* assays, Rab8a KO B cells showed diminished maturation into CD138^+^ antibody-producing cells, in agreement with the RNAseq data showing decreased CD138 expression. After immunisation we did not observe increased numbers of germinal centre cells, but the was a slight, although insignificant upregulation in the numbers of antibody-secreting cells (Fig. S2D-E).

Notably, we also found that Rab8a was specifically phosphorylated upon BCR, but, unlike in macrophages (*Luo et al*., 2014), not TLR stimulation (Fig. 5D). Rab8a has been reported to be phosphorylated by LRRK2, a tyrosine kinase protein highly expressed in immune cells, such as macrophages, neutrophils, and dendritic cells (*Wallings & Tansey*, 2019), and associated with Parkinson’s disease (*Steger et al*., 2016) and some autoimmune disorders, such as Crohn’s disease (*Barrett et al*., 2008; *Witoelar et al*., 2017). LRRK2 is required for the recruitment of Rab8a and Rab10 to the phagosomes in phagocytic cells (*Lee et al*., 2020) and it is upregulated following recognition of microbial structures (*Gardet et al*., 2010; *Hakimi et al*., 2011; *Kim et al*., 2012; *Moehle et al*., 2012). However, while the phosphorylation of Rab8a was partially downregulated in cells without functional LRRK2, inhibition or depletion of LRRK2 did not mirror the Rab8a KO phenotype in the *in vitro* CSR assay, suggesting that Rab8a function in B cells is independent of the phosphorylation by LRRK2 (Fig. 5E and Fig. S5).

To understand molecular mechanisms responsible for observed phenotypes we analysed BCR signalling in WT and Rab8a-deficient B cells. We observed no differences in proximal tyrosine kinase signalling and calcium signalling, but we found alterations in the downstream interactors (Fig. 7). Our data suggested that the kinetics of PI3K activation was dysregulated in Rab8a KO cells, and the levels of pERK1/2 and pAKT were decreased. Indeed, Rab8 has been shown to interact with PI3K in macrophages to regulate TLR signalling, inflammation and polarisation via the AKT/mTOR pathway (*Luo et al*., 2014, 2018; *Tong et al*., 2021; *Wall et al*., 2017, 2019). Thus, the enhanced CSR observed in Rab8a KO cells could be explained by the alterations in PI3K signalling, which in activated B cells is critical for CSR and the formation of GCs and plasma cells. Previous studies suggest that hyperactivation of PI3K signalling, either through activatory mutations, PTEN or FoxO1 deficiency, suppresses CSR, while promoting plasma cell differentiation (*Omori et al*., 2006; *Avery et al*., 2018; *Suzuki et al*., 2003). The regulation of CSR by PI3K is not completely understood as B cells express three isoforms of the class I PI3K catalytic subunit (p110α, p110δ, and p110γ) that can exert compensatory functions (*Donahue & Fruman*, 2004). In macrophages, Rab8a was shown to interact with p110γ (Luo *et al*., 2014, 2018; *Wall et al*., 2019) and PI3K catalytic subunit γ (PIK3CG) as well as Rab8a are both expressed in B cells and even at higher levels in human and mouse plasma cells (*ImmGen* and *plasmaCytomics* Datasets) highlighting the correlation between these two proteins. So, while we observed an abnormally early peak in PI3K phosphorylation, the levels were quickly settled lower than in the WT cells, which could ultimately lead to enhanced CSR (Fig. 5) and antibody secretion (Fig. 2 and 3) without increases in the plasma cell numbers (Fig. S2). The mechanisms how Rab8a could modulate p110γ in B cells and plasma cells warrants further examination. Partial inhibition of mTORC activity has also been suggested to augment CSR (*Limon et al*., 2014) and consistently with the decrease in pAkt, we detected a downshift in the phosphorylation of S6 protein suggesting a small decrease in Akt-mTORC1 signalling (Fig. 7D).

We also cannot rule out completely the effect of hypomorhic CD19, caused by CD19-cre system, in our Rab8a KO B cells, as this co-receptor is known to activate PI3K signalling downstream of BCR via binding to p85α PI3K regulatory subunit (*Wang et al*., 2002). CD19^-/-^ mice have abnormal B cell numbers and profound reduction in T-dependent immunisation-induced antibodies, but heterozygote complementation with human CD19 is able to restore the antibody levels (*Sato et al*., 1997). Also, this CD19-Cre model is widely used by the immunology community to conditionally ablate genes in B cells and it has, to best of our knowledge, not been linked to alterations in CSR.

Altogether, our data raises Rab8a as a potential regulator of antibody responses, plausibly via its linkage to PI3K and thereby to AKT/mTOR pathway leading to increased CSR and enhanced activity of antibody-secreting cells (Fig. S8). However, further studies are needed to verify the molecular mechanisms involved in this regulation.

## Materials and Methods

### Mice and cells

The CD19 Cre mouse colony (III) was backcrossed from a CD19 Cre RhoQ^flox/flox^ strain (*Burbage et al*., 2017), and the Rab8^flox/flox^ mouse colony was shared by Akihiro Harada (Osaka University, Japan) and have been previously described elsewhere (*Sato et al*., 2007). OT-II mice, expressing a transgenic TCR (Vα2/Vß5) specific for a peptide 323-339 of chicken ovalbumin presented by I-Ab (*Barnden et al*., 1998), were a kind gift from Maija Hollmén (University of Turku, Finland). Wildtype C57BL/6NCrl mice were purchased from the University of Turku Central Animal Laboratory and MD4 mice (C57BL/6-Tg(IghelMD4)4Ccg/J) from the Jackson Laboratory. C57BL/6-Lrrk2tm1.1Mjff/J mice (LRRK2 KO) developed by Michael J. Fox Foundation (The Jackson Laboratory, stock 016121) **(Baptista et al., 2013)** were a gift from Eleanor Coffey (Åbo Akademi, Finland). All strains were on a C57BL/6 background and maintained under specific-pathogen-free conditions. All experiments were done with age- (8-12 weeks), and sex-matched animals and WT littermate controls (CD19^WT/WT^, Rab8a^flox/flox^) were used whenever possible. When indicated, CD19 Cre mice (CD19^WT/Cre^, Rab8a^WT/WT^) or C57BL/6 mice were also used as control. All animal experiments were approved by the Ethical Committee for Animal Experimentation in Finland and adhered to the Finnish Act on Animal Experimentation (62/2006; animal license numbers: 7574/04.10.07/2014 KEK/2014-1407-Mattila, 10727/2018). A20 mouse lymphoma cells stably expressing a hen egg lysozyme (HEL)-specific IgM BCR (D1.3) (*Williams et al*., 1994) were a kind gift from Prof Facundo Batista (the Ragon Institute of MGH, MIT and Harvard, USA). Cells were maintained in complete RPMI [cRPMI; RPMI 1640 with 2.05 mM L-glutamine supplemented with 10% fetal calf serum (FCS), 50 μM β-mercaptoethanol, 4 mM L-glutamine, 10 mM HEPES and 100 U/ml penicillin/streptomycin].

### Immunisations

At the age of 8-10 weeks, groups of WT (CD19^WT/WT^ Rab8a^flox/flox^) and Rab8a^-/-^ (CD19^WT/Cre^ Rab8a^flox/flox^) females were immunised with NP40-FICOLL (F-1420, Biosearch Technologies) or NP-LPS (N-5065, Biosearch Technologies) for T-independent (TI) immunisation, or NP31-KLH (N-5060, Biosearch Technologies) for T-dependent (TD) immunisation. Each mouse received 50 μg of NP40-FICOLL in 150 μl of PBS, 25 μg of NP-LPS in 150 μl of PBS or 50 μl of NP31-KLH in 150 μl of Imject Alum-PBS (Thermo Fisher, #77161) by intraperitoneal injection. Blood (~100 μl) was sampled from lateral saphenous veins on day −1 (preimmunisation) and every week after immunisation on days +7, +14, +21, and +28 for both FICOLL and KLH cohorts. For the LPS cohort, blood was sampled on days −1, +3, +6, +9 and +12. Secondary immunisation of the NP-KLH cohort (50 μg in 150 μl of PBS) was performed on day +87 (0) and blood was sampled on days +85 (−2), +91 (+4), +95 (+8), and +102 (+15). Coagulated blood was spun at +4 °C/2500 rpm for 10 min, and serum was collected and stored at −20 °C.

### B and T Cell Isolation

Splenic B cells or T cells were Isolated with EasySepTM Mouse B Cell Isolation Kit (#19854, StemCell Technologies) or EasySepTM Mouse T Cell Isolation Kit (#19851, StemCell Technologies) according to the manufacturer’s instructions and let to recover in cRPMI (10% FCS, 20 mM HEPES, 50 μM β-mercaptoethanol, 50 U/ml potassium penicillin and 50 μg/ml streptomycin sulfate) in an incubator at +37 °C and 5% CO_2_ for 1 h before every experiment.

### Transfections

A20 D1.3 cells were transfected as previously described (*Šuštar et al*., 2018). Briefly, 3 million cells were resuspended in 180 μl of transfection buffer (5 mM KCl, 15 mM MgCl_2_, 15 mM HEPES, 50 mM sodium succinate, 180 mM Na_2_HPO_4_/ NaH_2_PO_4_ pH 7.2) containing 3 μg of plasmid and electroporated using AMAXA electroporation machine (program X-005, Biosystem) in 0.2 cm gap electroporation cuvettes. Cells were then transferred to 3 ml of cRPMI to recover overnight. The Rab8a-Citrine plasmid was a kind gift from Prof. Daniel Abankwa (University of Luxembourg).

### Western blot

Isolated splenic B cells were starved for 20 min in plain RPMI, and 2.5 × 10^6^ cells in 100 μl of plain RPMI were stimulated in duplicates with 10 μg/ml of F(ab)_2_ goat anti-mouse IgM antibodies either in solution or bound to the culture dish surface, for 5, 15 and 30 min. After activation, B cells were instantly lysed with RIPA buffer (sc-24948, Santa Cruz) with protease inhibitors (1X) and phosphatase (2 mM PMSF, 1 mM sodium orthovanadate, 10 mM NaF) inhibitors. Samples were incubated on ice for 30 minutes and centrifuged at 10.000 g for 20 min at 4 °C. Clear supernatants were transferred to a clean tube and quantified using DC Protein Assay (5000112, BioRad). Alternatively, primary B cells were activated as above and then lysed with 4X Laemmli buffer with beta-mercaptoethanol (BioRad) to a final concentration of 1X. Samples were sonicated for 5 minutes and boiled at 95 °C for 5 min. Unless specified otherwise, lysates (10-15 μg or 20-30 μl) were run on 10% polyacrylamide gels and transferred to PVDF membranes. For the Rab8 pT72 blots, nitrocellulose membranes were used. Membranes were blocked with 5% milk in TBS (TBS, pH 7.4) for 1 h and incubated with primary antibodies in 5% BSA in TBST (TBS, 0.05% Tween-20) O/N at 4 °C. Secondary antibody incubations (1:20.000) were done for 1 h at RT in 5% milk in TBST using HRP-conjugated secondary antibodies. Washing steps were done in 10 ml of TBST for 5 × 5 min. The following antibodies were used: anti-Rab8 (610844, BD Biosciences), anti-Rab8a (55296-1-AP, Proteintech; ab188574, Abcam; 6975, CST), anti-Rab8a phospho T72 (ab230260, Abcam), anti-beta-tubulin (66240-1-Ig, Proteintech), anti-beta Actin HRP (ab49900, Abcam), anti-phospho-Syk (2701, CST), anti-Syk (13198, CST), anti-phospho-Lyn (2731, CST), anti-Lyn (2796, CST), anti-phospho-AKT (Ser473) (4058, CST), anti-phospho ERK1/2 (9101, CST), anti-S6 (2217, CST), anti-phospho-S6 (4858, CST), and anti-GAPDH (60004-1-Ig, Proteintech). Membranes were scanned with ChemiDoc MP Imaging System (Bio-Rad) after the addition of Immobilon Western Chemiluminescent HRP Substrate (WBKLS0500, Millipore).

### Measurement of BCR internalisation by flow cytometry

To study the BCR internalisation upon activation with surrogate antigen, primary B cells (10^7^/ml) were stained for 5 min on ice with 10 μg/ml of biotinylated anti-IgM (Southern Biotech, 1021-08) and washed with Imaging Buffer. Cells (10^5^/well, 96-wp) were then incubated at 37°C and 5% CO2 for 45, 30, 15 and 5 min. As a control (time 0), samples were always kept on ice. After incubation, cells were stained with 1:1000 Alexa Fluor^®^ 633 streptavidin (Life Technologies, S-21375) in PBS on ice for 20 min. Samples were then washed with cold PBS and analysed. The internalisation rate for the biotinylated anti-IgM samples was calculated as described in Hernández-Pérez and Mattila, 2022. To study the constitutive BCR internalisation (in the absent of BCR activation), primary B cells (10^5^/well, 96-wp) were incubated at 37°C and 5% CO2 for 45, 30, 15 and 5 min in the presence or absence of 10 μg/ml of brefeldin A (BFA). As a control (time 0), samples were always kept on ice. After incubation, cells were stained with 1:500 Alexa Fluor^®^ 647 donkey anti-mouse IgM in PBS on ice for 20 min. Samples were then washed with cold PBS and analysed. The internalisation rate was calculated comparing the staining at each time point in the absence or presence of BFA. A BD LSR Fortessa analyser equipped with four lasers (405, 488, 561, and 640 nm) was used. Data were analysed using FlowJo v10 (Tree Star).

### Immunophenotyping

Immunophenotyping was done as previously described (*Sarapulov et al*., 2020). Bone marrow cells were isolated by flushing the buffer through mouse femoral and tibial bones, and splenocytes were isolated by mashing the spleen. All cells were isolated in Isolation Buffer (PBS, 2% FCS, 1 mM EDTA) and filtered through 70 μm nylon cell strainers. All subsequent steps were done in flow cytometry buffer I (PBS, 1% BSA). Fc-block was done with 0.5 μl of anti-mouse CD16/32 antibodies in 70 μl of flow cytometry buffer I for 10 min, and cells were stained for 30 min. Washings were done three times in 150 μl of flow cytometry buffer I. All steps were carried out on ice in U-bottom 96-well plates at a cell density of 0.25–0.5 × 106/well. Before the acquisition, cells were resuspended in 130 μl of flow cytometry buffer II (PBS, 2.5% FCS). Samples were acquired on BD LSR Fortessa, equipped with four laser lines (405, 488, 561, and 640 nm). The compensation matrix was calculated and applied to samples either in BD FACSDivaTM software (BD Biosciences) or in FlowJo (Tree Star, Inc) based on fluorescence of conjugated antibodies using compensation beads (Thermo Fisher, 01-1111-41). FMO (fluorescence minus one) controls were used to assist gating. Data were analysed with FlowJo software.

### B cell proliferation, class-switch recombination and antibody secretion

B cells purified from WT or KO spleens labelled with 5 μM of Cell Trace Violet (CTV) for 20 min at 37 °C in RPMI. Labelled B cells (10^5^ per well) were incubated with LPS (5 μg/ml), CD40L (150 ng/ml), IL-4 (5 ng/ml), IFN-γ (10 ng/ml), F(ab’)_2_ anti-IgM (10 μg/ml), CpG ODN (10 μg/ml), or a combination of those, in a 96 well U-bottom plate for 3 days (CSR and proliferation) or 5 days (antibody secretion). Cells were washed, blocked with Fc-block and stained as described in (*Sarapulov et al., 2020*). Samples were washed one more time and acquired on BD LSR Fortessa. B cell proliferation was analysed using ModFit. Class-switch was analysed using FlowJo. After 5 days, supernatants were transferred to a new plate for antibody quantification by ELISA.

### Immunohistochemistry

For immunisation experiments, spleens from non-immunised (alum) or NP-KLH-immunised animals were collected 9 days after immunisation. Spleens were embedded in O.C.T. compound and snap-frozen in chilled isopentane. 10 μm longitudinal sections were cut with a cryostat by the Histology core facility (Institute of Biomedicine, University of Turku, Finland). Sections were dried and fixed in 4% paraformaldehyde (PFA) for 15 minutes, followed by blocking with IF blocking buffer (10% FCS in PBS) for 1 h at RT. Sections were stained using a three-colour protocol with 1:100 of each of the following antibodies in IF blocking buffer at 4 °C O/N: Alexa Fluor 488^®^ anti-mouse GL-7 (GL-7; Biolegend, #144612), PE anti-mouse CD3 (145-2C11; Biolegend, #100308) and Alexa Fluor 647^®^ anti-mouse B220 (RA3-6B2; Biolegend, #103229). After washing 3 times with PBS, samples were mounted using Prolong Gold with DAPI (Thermo Fisher, #P36941). Sections were imaged using a Pannoramic Midi fluorescence slide scanner (3DHistech) equipped with a 20X objective and 4 filter sets (DAPI, FITC, Rhodamine/TRITC, Cy5). Tiled sections were exported to TIFF format and analysed with ImageJ.

### ELISpot

NP-specific antibody-secreting cells (ASCs) were measured by ELISpot. ELISpot plates (Mabtech, #3654-TP-10) were pre-wet with 15 μl/well of freshly prepared 35% ethanol for < 1 min at RT. Plates were washed twice with 150 μl/well of sterile water, and wells were coated O/N with 25 μg/ml NP24-BSA in PBS at 4 °C. The next day, plates were washed with PBS and blocked with 200 μl/well of cRPMI for 2 h at RT. Splenocytes were isolated, resuspended in cRPMI and seeded in the wells performing serial dilutions (6, 3 and 1.5 × 105 cells/well). Cells were incubated at 37 °C, 5% CO_2_ for 18 hours. Cells were washed 5 times with 150 μl/well washing buffer (PBS, 0.05% Tween-20). Biotin-conjugated detection antibodies (2 μg/ml) in 100 μl of blocking buffer (1% BSA in PBS) were added for 2 h at RT followed by 100 μl streptavidin-HRP (1:000) in blocking buffer for 1 h at room temperature (RT). Filtered (0.45 μm pore) TMB (Matbtech, #3651-10) was then added to the plates for 5 minutes, and wells were extensively washed under running mQ water. ELIPspot plates were imaged using a Zeiss Axio-Zoom.V16 optic microscope.

### Eα peptide presentation

Antigen presentation was measured using the Eα peptide system (*Viret & Janeway*, 2000). For coating, 200 nm Dragon Green Streptavidin beads (Bangs Laboratories, #CFDG001, Lot 13743) were used. Beads were washed with 2% FCS in PBS, sonicated to disrupt bead:bead aggregates (Bioruptor^®^: HIGH, 5 cycles, 30” on/off) and coated with different ratios of biotinylated anti-IgM (Southern Biotech, #1021-08) and biotinylated Eα52-68 peptide (Biotin-GSGFAKFASFEAQGALANIAVDKA-COOH) for 1 h at 37 °C. Beads coated with biotinylated anti-IgM alone were used as a control. Cells and beads were incubated for 30 min at 37 °C to allow binding and internalisation, washed, and incubated again at 37 °C for 4 hours. After 4 hours, samples were transferred on ice, washed, blocked with Fc-block and stained with anti-Eα-MHC-II antibody (Invitrogen, #14-5741-85) for 30 minutes. Then, samples were washed and stained with Alexa Fluor^®^ 633 goat anti-mouse IgG2b (1:500) for another 30 minutes. Samples were washed one more time and acquired on BD LSR Fortessa.

### OT-II proliferation

For coating, 100 nm Streptavidin beads (Bangs Laboratories, #CP01000, Lot 14817 and CP01003 Lot 10045) were used. Beads were prepared as described for Eα peptide presentation and coated with different ratios of biotinylated anti-IgM (Southern Biotech, #1021-08) and biotinylated ovalbumin (OVA) (produced in-house) for 1h at 37 °C. Beads coated with biotinylated anti-IgM alone were used as a control. B cells purified from WT or KO spleens labelled with 1 μM CFSE and T cells purified from OT-II spleens were labelled with 5 μM Cell Trace Violet (CTV) for 20 min at 37 °C in RPMI. Labelled B cells were incubated with the beads for 30 min at 37 °C, washed, and incubated with the labelled T cells (1:1) in a 96 well U-bottom plate for 72 hours. After 3 days, supernatants were transferred to a new plate for cytokine quantification in ELISA and cells were washed, blocked with Fc-block and stained with LiveDead APCe780 (1:1000; Thermo Fisher, #65-0865-14), APC anti-CD19 (1:200; BioLegend, #392503) and PE anti-CD3 (1:200; Biolegend, #100308). Samples were washed one more time and acquired on BD LSR Fortessa. B and T proliferation was analysed using the Proliferation Module in FlowJo.

### ELISA

IL-6 secretion was quantified using an ELISA MAX Deluxe Set Mouse IL-6 according to the manufacturer’s instructions (431304, Biolegend). Total and NP-specific antibody levels and IL-2 secretion were measured by ELISA on half-area 96-well plates (Greiner Bio-One, 675061) as previously described (*Hernández-Pérez et al*., 2020; *Sarapulov et al*., 2020). Wells were coated overnight at 4 °C with capture antibodies (2 μg/ml) or NP-conjugated carrier proteins (50 μg/ml; NP(19)-BSA for high affinity or NP(>20)-BSA for all affinities (N-5050L, N-5050H, Biosearch Technologies)) in 25 μl PBS. Non-specific binding sites were blocked for 1-2 h in 150 μl of blocking buffer (PBS, 1% BSA) and 50 μl of serum samples or supernatants diluted in blocking buffer were added for overnight incubation at +4 °C. Results reported correspond to the following serum dilutions: 1:50.000 for total or isotype-specific IgM and IgG2b, 1:75.000 for IgG1 and 1:25.000 for IgG2c and IgG3 antibodies. Biotin-conjugated detection antibodies (2 μg/ml) in 50 μl of blocking buffer were added for 1 h followed by 50 μl ExtrAvidin-Alkaline phosphatase (E2636, Sigma-Aldrich, 1:5.000 dilution) in blocking buffer for 1 h at RT. In between all incubation steps, plates were washed with 150 μl washing buffer (PBS, 0.05% Tween-20) either three times for the steps before sample addition or six times after the addition of the mouse sera. The final wash was completed by washing two times with 150 μl of MQ water. Finally, 50 μl of alkaline phosphatase-substrate, SIGMAFAST p-nitrophenyl phosphate (N2770, Sigma-Aldrich) solution (0.5 – 1 mg/ml) was added and OD was measured at 405 nm.

All ELISA samples were run in duplicates, OD values were averaged, and blank background was subtracted. Absolute concentrations of total antibody levels were extrapolated from calibration curves prepared by serial dilution of mouse IgM or subclasses of IgG from C57BL/6 immunoglobulin panel. Relative NP-specific antibody levels were extrapolated from reference curves prepared by serial dilution of pooled serum from day 14 after NP-KLH immunisation, in which the highest dilution step received an arbitrary unit value.

### B cell activation and visualisation of antigen vesicles by immunofluorescence microscopy

Twelve-well PTFE diagnostic slides (Thermo Fisher, #10028210) were coated with 4 μg/ml fibronectin in PBS. A20 D1.3 or primary B cells isolated from MD4 mice spleens were labelled on ice for 10 minutes with 10 μg/ml of Alexa Fluor^®^ 647 donkey anti-mouse IgM (#715-605-140, Jackson ImmunoResearch) or RRx goat anti-mouse IgM (#115-295-205, Jackson ImmunoResearch), washed with PBS to remove excess unbound antigen and resuspended in Imaging Buffer (PBS, 10% FCS). After washing, cells were seeded on the fibronectin-coated wells and incubated at 37 °C to trigger activation. Then, cells were fixed with 4% PFA for 10 min at RT and blocked/permeabilised for 20 min at RT (5% donkey serum with 0.3% Triton X-100 in PBS). After blocking, samples were stained with primary antibodies for 1h at RT or 4 °C O/N in staining buffer (1% BSA, 0.3% Triton X100 in PBS), followed by washes with PBS and incubation with the fluorescently-labelled secondary antibodies for 30 min at RT in PBS. Samples were mounted using FluoroMount-G containing DAPI (Thermo Fisher, #00495952). Images were acquired on a 3i CSU-W1 Marianas spinning disk confocal microscope (Intelligent Imaging Innovations) equipped with a 63× Zeiss Plan-Apochromat objective (NA 1.4) and a Hamamatsu sCMOS Orca Flash4.0 camera (2048 × 2048 pixels, 1 × 1 binning).

All SDCM images were deconvolved with Huygens Essential version 16.10 (Scientific Volume Imaging, The Netherlands, http://svi.nl), using the CMLE algorithm, with Signal to Noise Ratio of 20 and 40 iterations. Colocalisation analysis was done on ImageJ using the Colocalisation Threshold plugin. As a control for random colocalisation, the antigen channel was rotated 90° and colocalisation was measured using the plugin.

### Synapse formation

Twelve-well PTFE diagnostic slides (Thermo Fisher, #10028210) were coated with 5 μg/ml donkey anti-mouse IgM (#715-005-020, JIR) or Cell-Tak (Corning) in PBS at RT for at least 1 hour. Primary B cells isolated from WT and KO spleens were resuspended in RPMI (106 cells/ml) and labelled, or not, with 0.5 μM CTV (Thermo Fischer, C34557). Labelled and unlabelled cells were mixed (1:1 ratio) and seeded at a density of 80.000 cells/well. Dye-switched experiments were performed systematically. Cells were activated for 15 min (+37 °C, 5% CO_2_), fixed in 4% PFA for 10 min at RT, and permeabilised/blocked (5% donkey serum with 0.3% Triton X100 in PBS) for 20 min at RT. Cells were stained with Alexa Fluor^®^ 488 anti-phospho-PLCγ2 (1:100; BD Biosciences, 558507) or Alexa Fluor^®^ 488 anti-phospho-Btk (1:100; BD Biosciences, 564847) and Acti-Stain 555 Phalloidin (1:150; Cytoskeleton Inc., PHDH1-A) for 1 hour in staining buffer. Samples were mounted in FluoroMount-G (Thermo Fisher). Images were acquired on a 3i CSU-W1 Marianas spinning disk confocal microscope (Intelligent Imaging Innovations) equipped with a 63× Zeiss Plan-Apochromat objective (NA 1.4) and a Photometrics Prime BSI sCMOS camera (2048×2048 pixels, 1×1 binning). Cells were visualised at the contact plane, and 5–10 fields of view per sample were acquired. Images of F-actin, pBtk, and pSyk were processed with ImageJ using the CTV channel to discriminate between WT or Rab8a KO cells. Spreading area (determined on the phalloidin channel) and mean fluorescence intensity of pBtk or pSyk staining per cell were analysed (> 100 cells per condition per experiment, at least n = 4 independent experiments).

### Intracellular Ca ^2+^ Flux

Splenic B cells were resuspended at a concentration of 5 × 106 cell/ml in RPMI supplemented with 20 mM HEPES and 2.5% FCS and loaded with 1 μM Fluo-4 (Thermo Fisher, #F14201) and 3 μM Fura Red (Thermo Fisher, #F3021) for 45 min (+37 °C, 5% CO_2_) with occasional mixing (*Sarapulov et al*., 2020). After washing with cRPMI, cells were resuspended at 2.5 × 106 cells/ml in PBS supplemented with 20 mM HEPES, 5 mM glucose, 0.025% BSA, 1 mM CaCl_2_, 0.25 mM sulfinpyrazone (Sigma-Aldrich, #S9509), and 2.5% FCS. A 30-second baseline reading was recorded for each sample before addition of F(ab’)_2_ goat anti-mouse IgM (5-10 μg/ml). Calcium flux was monitored for a total time of 5 minutes. Samples were acquired on BD LSR Fortessa using Tube mode, and data (n = 3 independent experiments) were analysed with FlowJo software and presented as the ratio-metric measurement of Fluo-4/Fura Red geometric mean (GeoMean) intensity levels.

### Transwell migration / Chemotaxis assay

Chemotaxis assays were carried out using 24-well Transwell chambers with 5-μm pore size polycarbonate membranes (Sigma, #CLS3421-48EA). B cells (107 cells/ml) were labelled with 0.1 μM CFSE or 0.5 μM Cell Trace Violet (CTV) for 20 minutes in the incubator (37 °C, 5% CO_2_). After incubation, cells were washed twice with cRPMI and resuspended in chemotaxis medium (0.5% BSA, 10 mM HEPES in RPMI) to a concentration of 107 cells/ml. WT and KO cells were mixed 1:1 ratio, and colour-switched experiments were systematically performed. The chemotaxis medium with or without CXCL13 (1 μg/ml; PeproTech #250-24) or CXCL12/SDF-1α (100 ng/mL; Bio-Techne, #460-SD-010) was added to the wells of the receiving plate, and 100 μL cell suspension (106 cells/sample) was placed in the transwell insert. Samples were incubated for 4 hours at 37 °C in humidified air with 5% CO_2_. After 4 hours, transwell inserts were removed, and the migrated cells in the receiving wells were recovered. Recovered cells were stained with eBioscience™ Fixable Viability Dye eFluor™ 780 (FVD; Thermo Fischer) and Alexa Fluor^®^ 647 anti-mouse CD19 in PBS-2% FCS containing Fc-block for 30 minutes on ice. After 2 washes in PBS-2% FCS, cells were resuspended in 100 μl of PBS. A known volume of each sample, as well as the initial cell mix, was acquired on BD LSR Fortessa equipped with an HTS. After excluding cell debris and gating on singlets and CD19+ FVD-cells, the percentage of migrating cells (migration index) was calculated by dividing the total number of migrated cells in each well by the total number of input cells determined by FACS x 100.

### Treatment with LRRK2 inhibitors

For Western blot, A20 D1.3 cells were pre-treated with GNE-7915 or GSK2578215A (MedChem Express, HY-18163 and HY-13237) for 1 or 2 hours followed by stimulation with 10 μg/ml of F(ab)_2_ goat anti-mouse IgM antibodies as described above. For CSR experiments, primary B cells were isolated and labelled with Cell Trace Violet as described above and inhibitors (0.5, 1 and 5 μM) were added to the medium and kept there for the duration of the experiment (72 h).

### RNA isolation and RNAseq

RNA was extracted from a total of 10^6^ B cells purified from spleen or lymph nodes or 105 B cells sorted from the B:T co-cultures (72 h) using TRIsure™ (Bioline) following the manufacturer’s instructions. When using < 10^7^ cells per sample, RNase-free Glycogen Co-precipitant (Bioline) was added to aid RNA precipitation.For RNAseq, RNA samples were sent to Novegene (Cambridge, UK). After quality check (RIN > 8), messenger RNA was subjected to library construction and sequenced on the Illumina NovaSeq 6000 platform (PE150, 20 M read pairs per sample). Additional details can be found in Supplementary Materials and Methods.

### Statistical analysis and illustrations

Statistical significance was calculated using unpaired Student’s t-test assuming a normal distribution of the data unless otherwise stated in the figure legends. Statistical values are denoted as: *P < 0.05, **P < 0.01, ***P < 0.001, ****P < 0.0001. Graphs were created in GraphPad Prism 6/8, and illustrations were created with BioRender. Figure formatting was done on Inkscape 1.0.

## Author Contributions

Conceptualisation: SHP, PKM. Formal Analysis: SHP. Funding Acquisition: PKM. Investigation: SHP, AVS, MOB. Methodology: SHP, PKM. Project Administration: SHP, PKM. Resources: PKM, AH, EC. Supervision: SHP, PKM. Validation: SHP, PKM. Visualisation: SHP, AVS. Writing – Original Draft Preparation: SHP. Writing – Review & Editing: SHP, AVS, PKM.

## Acknowledgments

We thank Laura Grönfors, Amna Music and Anni Virtänen for technical assistance. We thank Marika Runsala and Dr Elina Kuokkanen for collecting preliminary data. We thank Dr Ana Martínez Riaño and Dr Marianne Burbage for help in troubleshooting the Eα peptide system, Dr Søren Degn for scientific advising, Dr Johan Peränen for generous sharing of reagents, Alex Barkoff for help with the ELISpot system, and Dr Virginia García de Yébenes and Dr María Isabel Yuseff for comments on our work. Microscopy and flow cytometry were performed at Turku Bioscience Cell Imaging and Cytometry (CIC), supported by Turku Bioimaging and Euro-Bioimaging. The research infrastructures were provided by Biocenter Finland and InFLAMES.

## Data Availability

All data is available upon request.

## Supplementary Information

- Supplementary Figures 1–8
- Supplementary Materials and Methods

**Figure S1:**
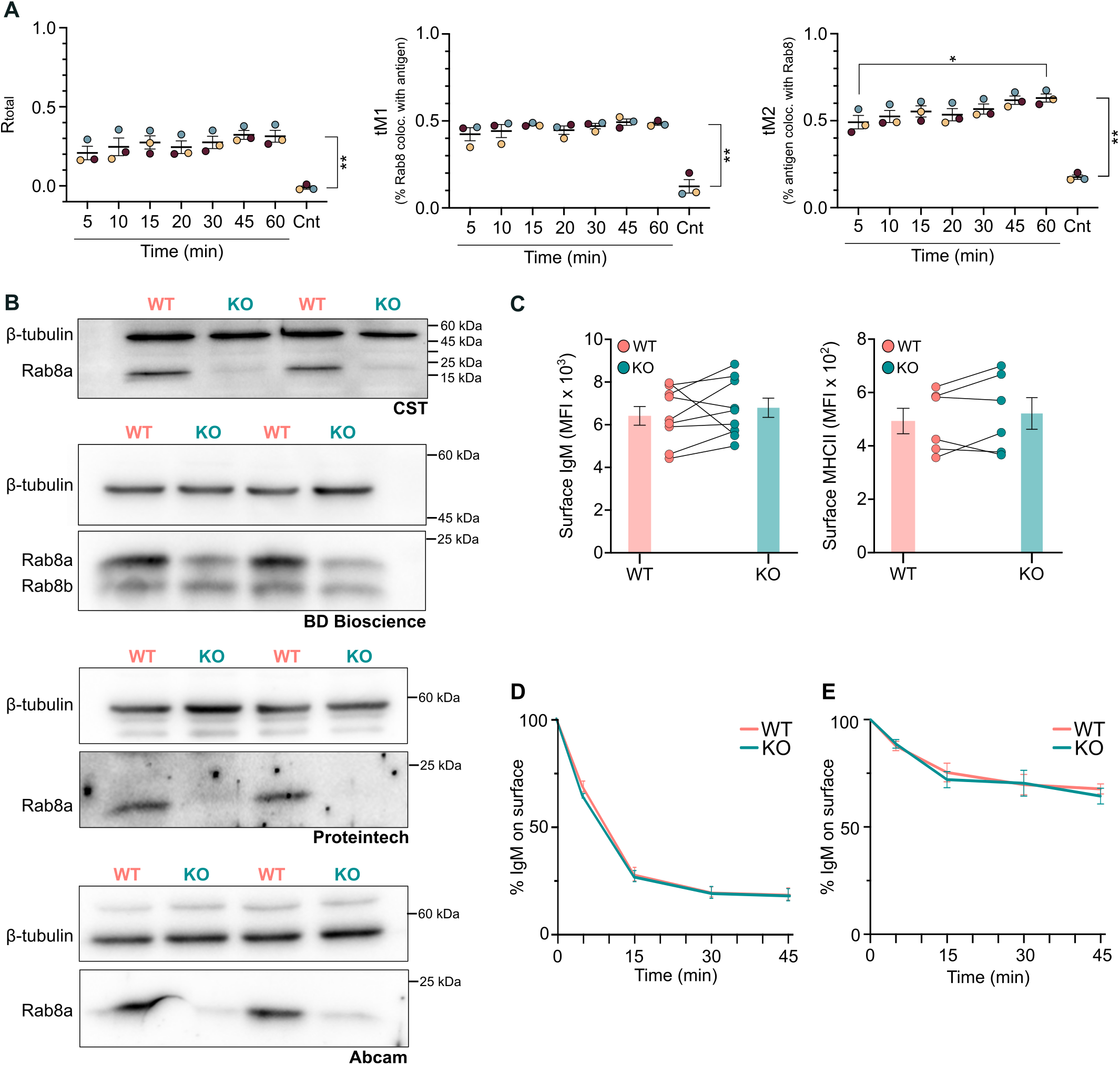
Rab8a shows constant colocalisation with internalised antigen and the surface levels of BCR (IgM) and MHCII, as well as BCR internalisation, are unaffected in Rab8a-KO mice. (A) Colocalisation of Rab8a with the antigen analysed in 3 independent experiments. Each dot represents the mean of one experiment (mean ± SEM; cells 30-80 per timepoint per experiment). As a negative control, to evaluate the random colocalisation, the Rab8a channel was rotated 90° right. Statistics: unpaired t-test. (B) Expression of Rab8a/b by Western blot in WT and Rab8a KO mice. Membranes were probed with anti-Rab8 antibodies from different companies (indicated below each blot). CST, Proteintech and Abcam antibodies recognise Rab8a, while BD Bioscience antibody recognise Rab8a and Rab8b. (C) Surface levels of IgM and MHCII analysed with flow cytometry. N = 6-9 experiments (mean ± SEM). (D) BCR internalisation after activation with anti-IgM (n = 3; mean ± SEM). (E) Constitutive BCR internalisation in the absence of activation (n = 3; mean ± SEM).

**Figure S2:**
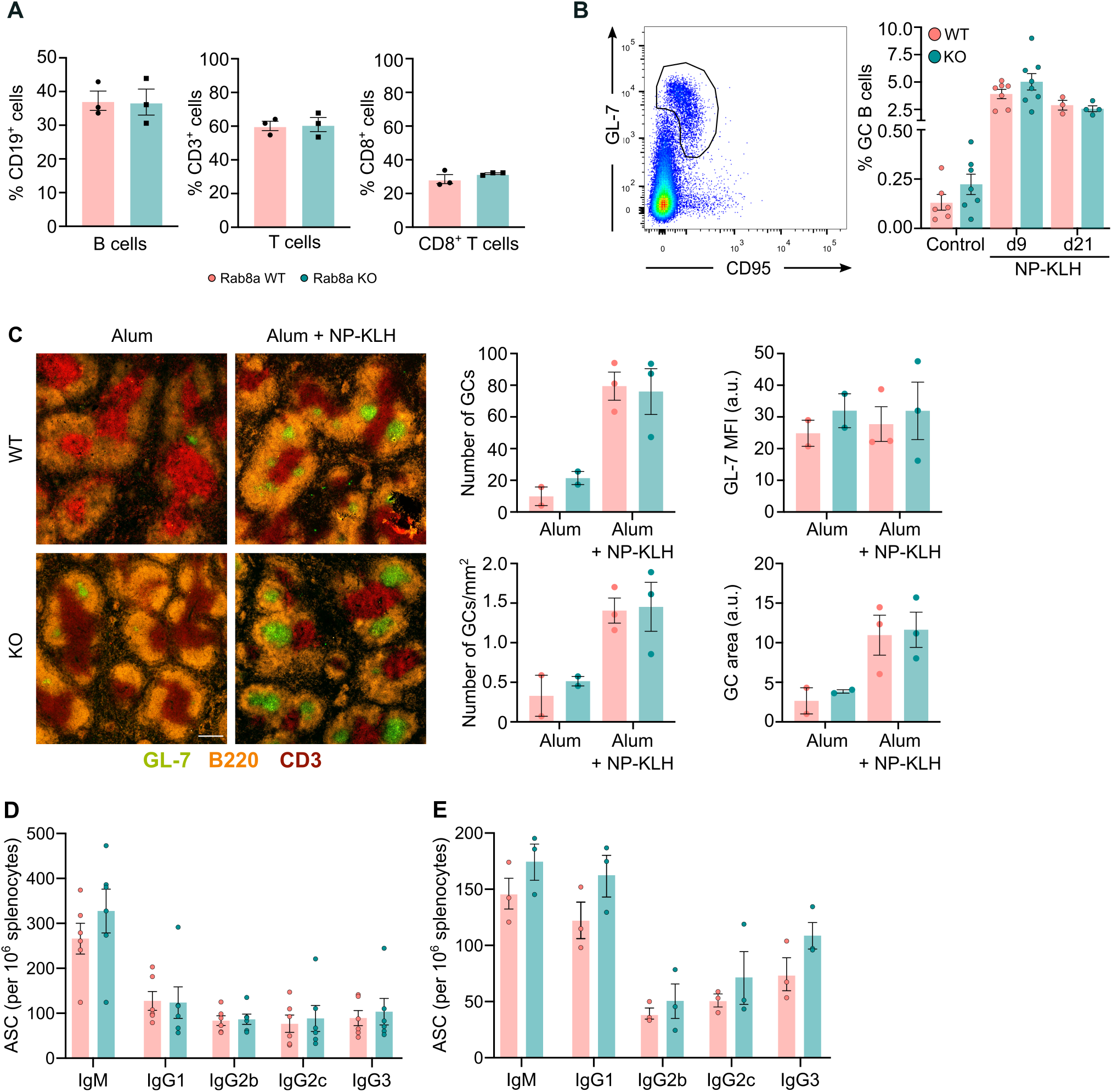
Rab8a KO mice have normal GC formation and normal numbers of antigen-secreting cells. **(A)** B and T cell populations in the lymph nodes from WT and Rab8a KO mice (n = 3 mice; mean ± SEM). **(B)** Splenocytes from WT or KO mouse were analysed by flow cytometry in non-immunised mice (control), or NP-KLH (in alum) immunised mice on day +9 and +21. Germinal centre B cells (CD19+ GL7+ CD95+) were gated as shown on the left plot. Data from n = 3-8 independent experiments (each dot represents one mouse) are shown (mean ± SEM). Paired t-test. **(C)** Spleens from control (alum) or NP-KLH + alum immunised WT or KO mice on day +9 analysed were snap-frozen, cut and stained for GCs (GL-7, in green). Sections were imaged using a Pannoramic slide scanner. The number of GCs (total number and normalised by mm2), the germinal centre size (area) and the intensity of the GL-7 staining are shown on the right (n = 3 independent experiments, each dot is the mean from one experiment; mean ± SEM) **(D)** Splenocytes from NP-KLH (in alum) immunised WT or KO mice on day +9 analysed by ELISpot. N = 3 independent mice pairs. Below the graph, representative images of the ELISpot plates. **(E)** Splenocytes from NP-FICOLL immunised WT or KO mice on day +9 analysed by ELISpot. N = 3 independent mice pairs.

**Figure S3:**
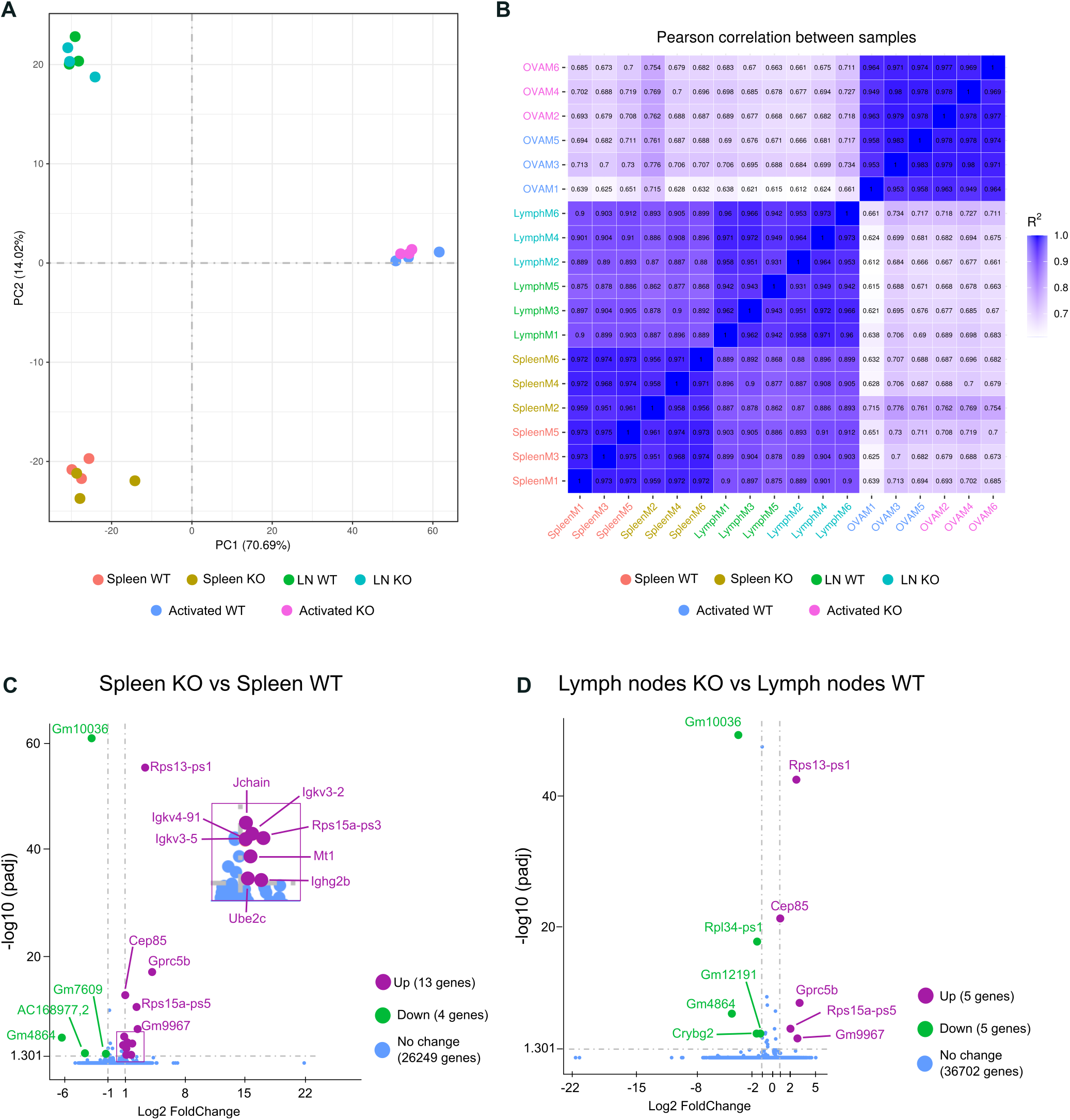
RNA sequencing. **(A)** Principal component analysis of the RNAseq samples. LN = lymph nodes (n = 3 mice per condition). **(B)** Pearson correlation between samples (spleen non activated; lymph nodes LN non activated or OVA activated). N = 3 mice per condition (M1, M3, M5 = WT; M2, M4, M6 = KO). **(C-D)** Volcano plot showing the DEGs in **(C)** resting spleen and **(D)** resting lymph nodes of WT and Rab8a KO mice.

**Figure S4:**
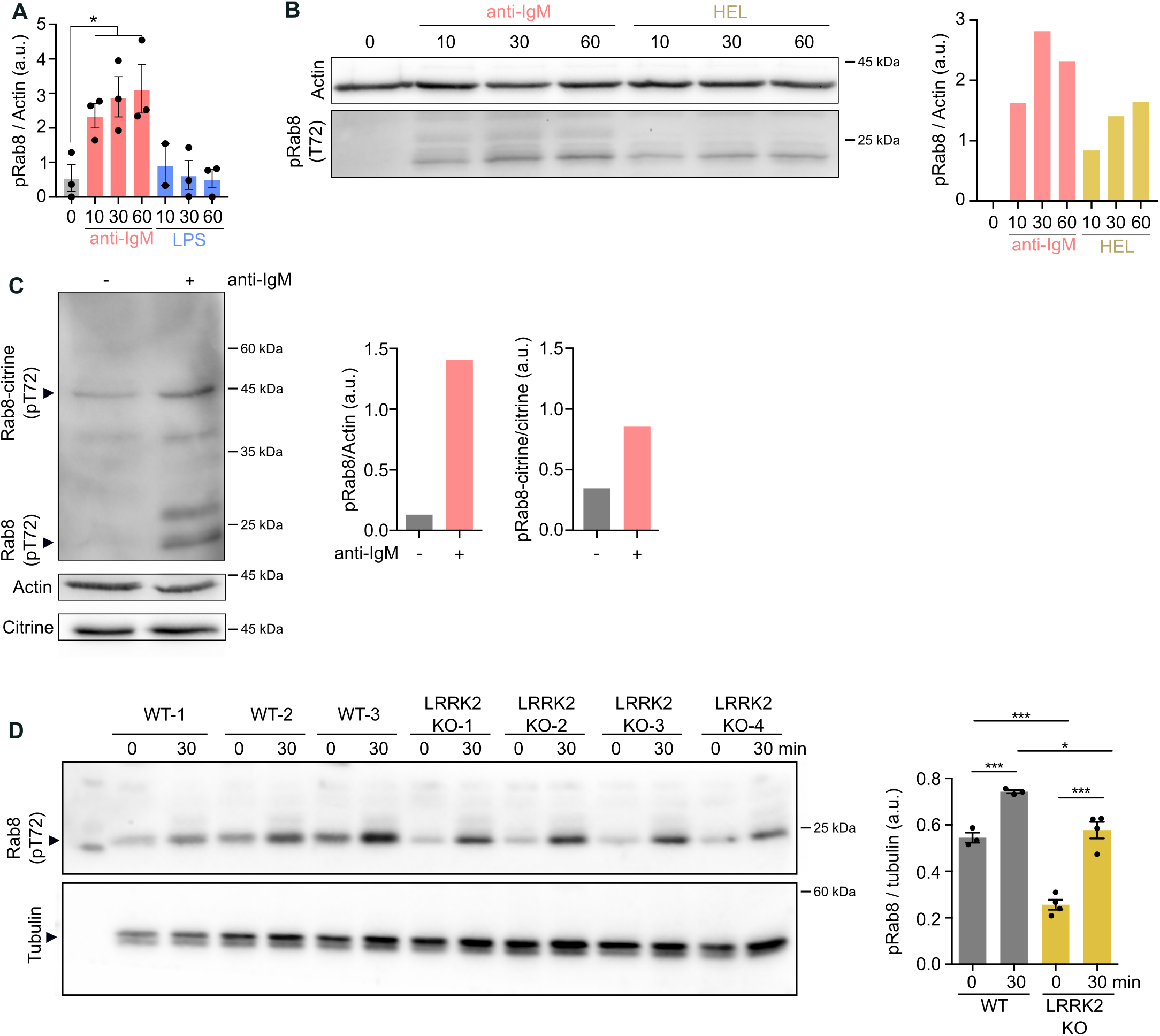
Rab8a phosphorylation in response to BCR activation and in LRRK2 KO cells. **(A)** Quantification of Fig. 5D. Mean ± SEM. Statistics: unpaired t-test.**(B)** Rab8 phosphorylation was studied by Western Blot in response to BCR stimulation either with surrogate antigen (F(ab’)_2_ anti-IgM) or HEL in A20 D1.3 cells (representative experiment). The levels of pRab8 were normalised to actin (loading control). **(C)** Rab8 phosphorylation in A20 D1.3 cells transfected with Rab8a-citrine analysed by WB (representative experiment). **(D)** Primary B cells isolated from WT or LRRK2 KO mice were activated (30 min) or not (0 min) with F(ab’)_2_ anti-IgM and blotted for anti-pT72 Rab8. Tubulin was used as a loading control. The right graph shows the quantification (mean ± SEM; n = 3 WT mice and 4 KO mice). Statistics: unpaired t-test.

**Figure S5:**
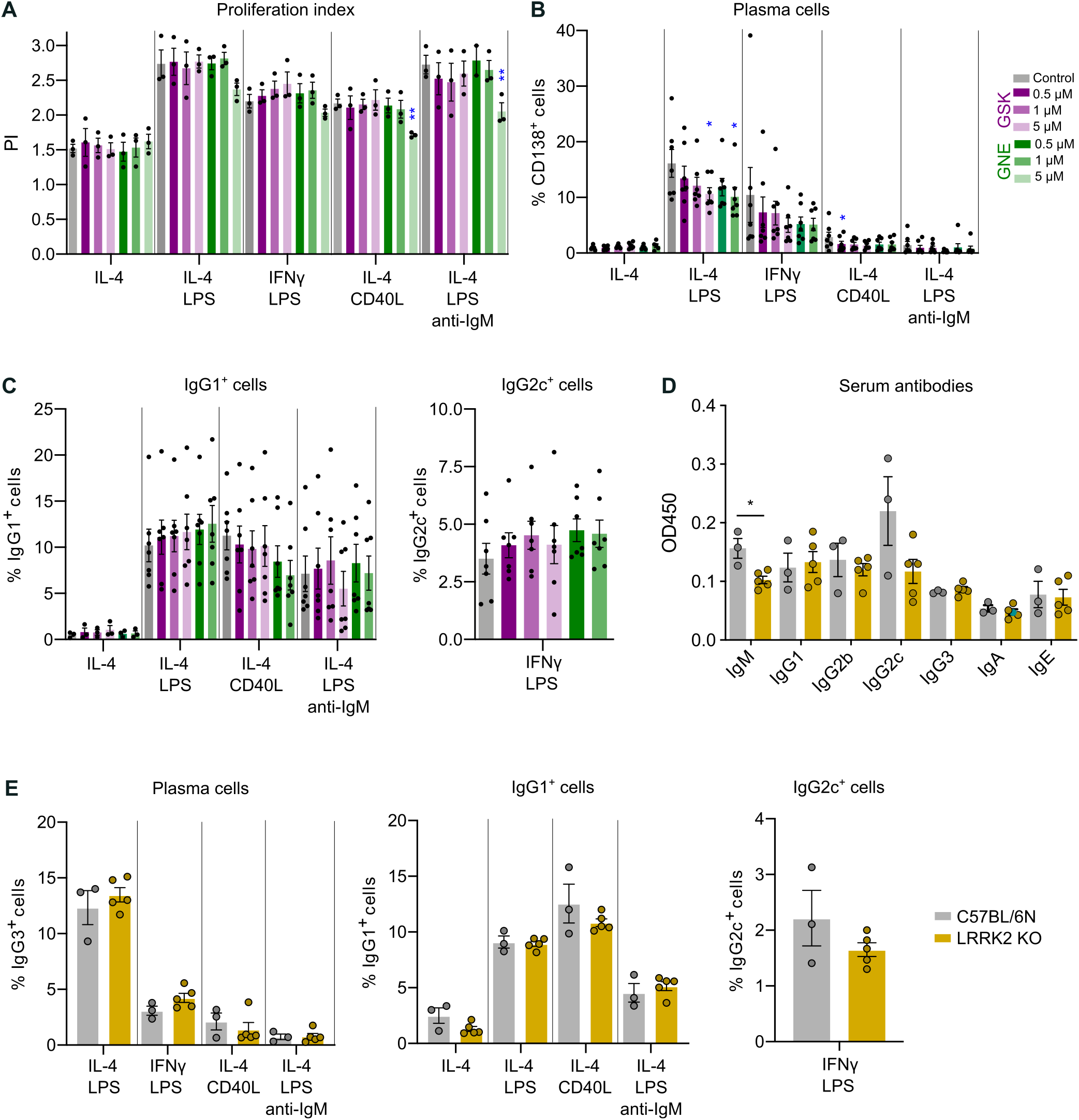
LRRK2 inhibition does not recapitulate the Rab8 KO phenotype. **(A-C)** Primary WT B cells were untreated or treated with 0.5,1 and 5 μM of GSK or GNE, and stimulated with the combinations shown in the graph. **(A)** Proliferation was analysed using the Proliferation module in FlowJo. Mean ± SEM (n = 3 independent experiments). Statistics: unpaired t-test. **(B)** Plasma cells were determined by flow cytometry gating on the CTVlow CD138+ cells and **(C)** class-switched cells were analysed gating on the CTVlow IgG1+ or IgG2c+ cells. Mean ± SEM (n = 7 independent experiments). **(D)** Basal levels of serum antibodies measured in the blood of LRRK2 KO mice (n = 5) compared to control C57BL/6 mice (n = 3). Mean ± SEM. Statistics: unpaired t-test.

**Figure S6:**
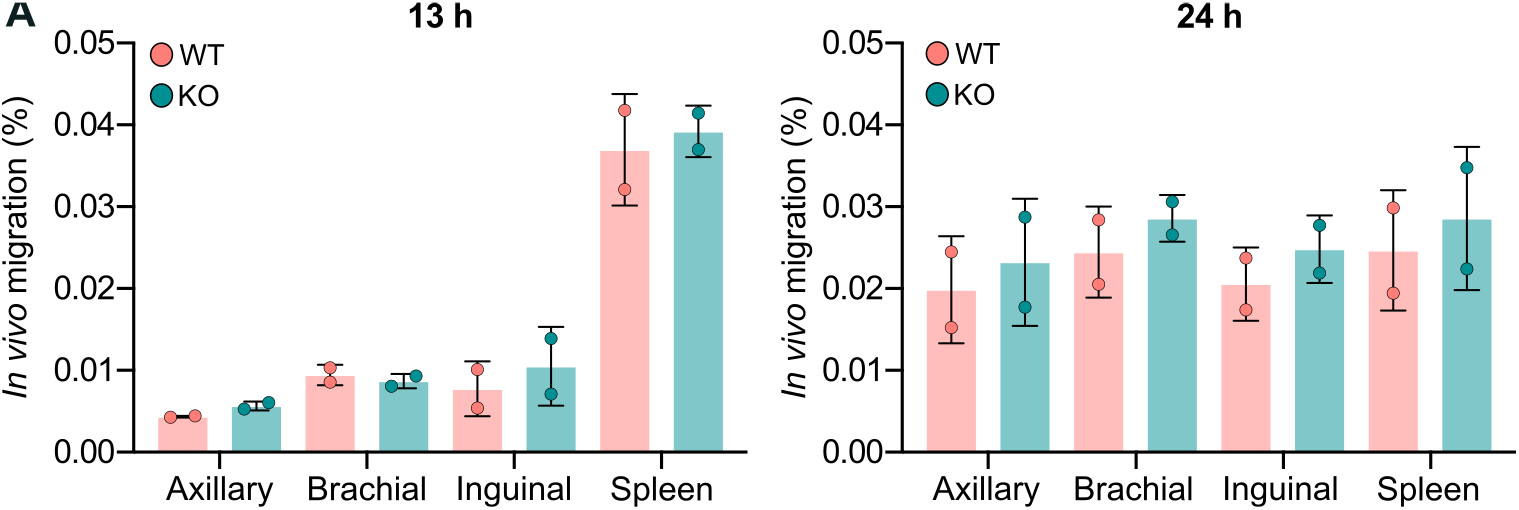
Rab8a does not affect *in vivo* cell migration *In vivo* migration. Primary mouse B cells from WT or Rab8 KO mice were labelled with CTV and CFSE and the cells (1:1) were injected in the tail vein of a recipient WT mouse. The mice were culled 13 or 24 h after the injection and the homing of B cells to different lymph nodes and spleen were analysed by flow cytometry. Two mice per time point were analysed.

**Figure S7:**
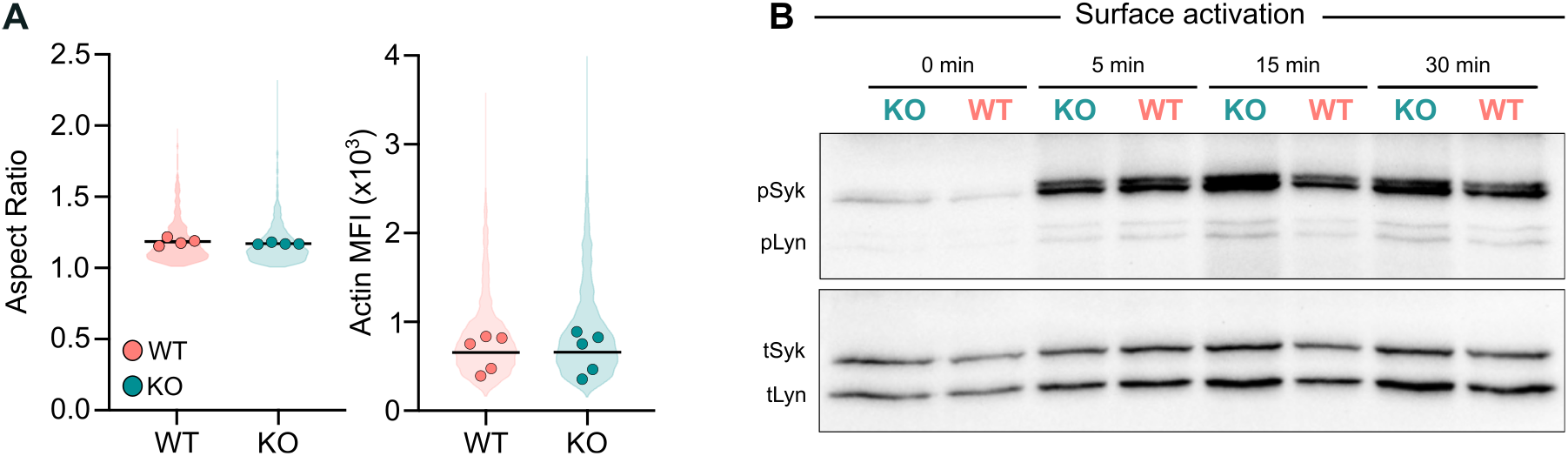
Spreading response and surface-bound activation. **(A)** Spreading on anti-IgM coated glass; related to Fig. 7A. B cells from WT or Rab8a KO mice were seeded on anti-IgM coated glass for 15 minutes, fixed and stained with phalloidin (actin). The spreading was analysed by measuring the Aspect Ratio (symmetry of the synapse) and the intensity of the actin staining. The violin plots represent the distribution of the population (all analysed cells, > 100 cells per experiment) and dots represent the mean of individual experiments (n = 4-5). **(B)** Proximal kinase signalling (pSyk and pLyn) in response to plate-bound F(ab’)_2_ anti-IgM by Western blot. Primary B cells were activated on a plate coated with 10 μg/ml of F(ab’)2 anti-IgM for 5,15 and 30 minutes. Statistics: paired t-test.

**Figure S8:**
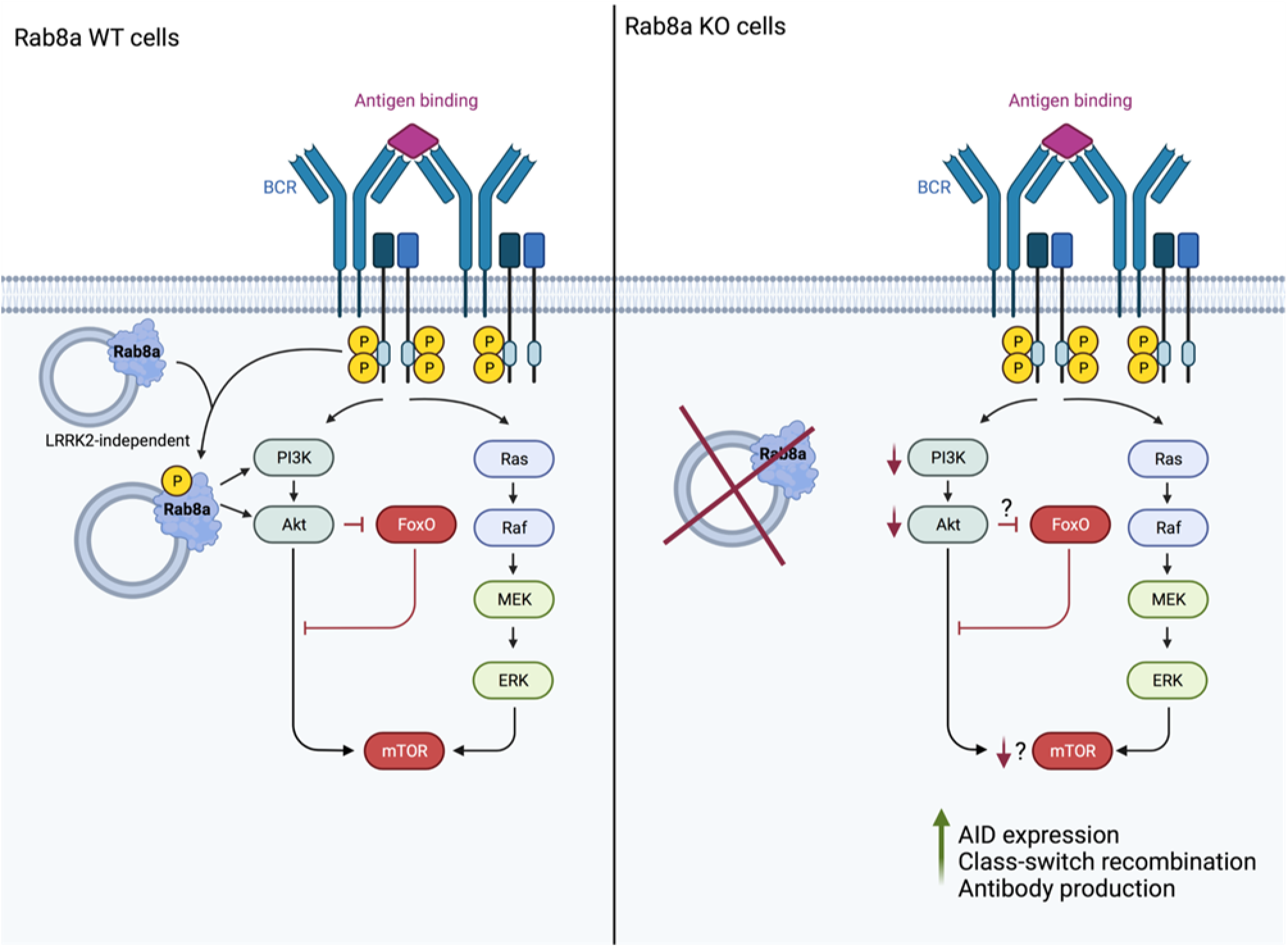
Loss of Rab8 alters the PI3K/AKT/mTOR signalling, leading to increased AID expression and CSR.

## Supplementary Materials and Methods

### RNAseq

#### Library Construction, Quality Control and Sequencing

Messenger RNA was purified from total RNA using poly-T oligo-attached magnetic beads. After fragmentation, the first strand cDNA was synthesised using random hexamer primers, followed by the second strand cDNA synthesis using either dUTP for directional library or dTTP for non-directional library. The library was checked with Qubit and real-time PCR for quantification and bioanalyzer for size distribution detection. Quantified libraries will be pooled and sequenced on Illumina platforms, according to effective library concentration and data amount.

#### Clustering and sequencing

The clustering of the index-coded samples was performed according to the manufacturer’s instructions. After cluster generation, the library preparations were sequenced on an Illumina platform and paired-end reads were generated.

### Data analysis

#### Quality control

Raw data (raw reads) of fastq format were firstly processed through in-house perl scripts. In this step, clean data (clean reads) were obtained by removing reads containing adapter, reads containing ploy-N and low-quality reads from raw data. At the same time, Q20, Q30 and GC content the clean data were calculated. All the downstream analyses were based on the clean data with high quality.

#### Reads mapping to the reference genome

Reference genome and gene model annotation files were downloaded from genome website directly. Index of the reference genome was built using Hisat2 v2.0.5 and paired-end clean reads were aligned to the reference genome using Hisat2 v2.0.5. We selected Hisat2 as the mapping tool for that Hisat2 can generate a database of splice junctions based on the gene model annotation file and thus a better mapping result than other non-splice mapping tools.

#### Quantification of gene expression level

featureCounts v1.5.0-p3 was used to count the reads numbers mapped to each gene. And then FPKM of each gene was calculated based on the length of the gene and reads count mapped to this gene. FPKM, expected number of Fragments Per Kilobase of transcript sequence per Millions base pairs sequenced, considers the effect of sequencing depth and gene length for the reads count at the same time, and is currently the most commonly used method for estimating gene expression levels.

#### Differential expression analysis

Differential expression analysis of two conditions/groups (two biological replicates per condition) was performed using the DESeq2 R package (1.20.0). DESeq2 provide statistical routines for determining differential expression in digital gene expression data using a model based on the negative binomial distribution. The resulting P-values were adjusted using the Benjamini and Hochberg’s approach for controlling the false discovery rate. Genes with an adjusted P-value <=0.05 found by DESeq2 were assigned as differentially expressed.

#### Enrichment analysis of differentially expressed genes

Gene Ontology (GO) enrichment analysis of differentially expressed genes was implemented by the clusterProfiler R package, in which gene length bias was corrected. GO terms with corrected Pvalue less than 0.05 were considered significantly enriched by differential expressed genes. KEGG is a database resource for understanding high-level functions and utilities of the biological system, such as the cell, the organism and the ecosystem, from molecular-level information, especially large-scale molecular datasets generated by genome sequencing and other high-through put experimental technologies (http://www.genome.jp/kegg/). We used clusterProfiler R package to test the statistical enrichment of differential expression genes in KEGG pathways. The Reactome database brings together the various reactions and biological pathways of human model species. Reactome pathways with corrected Pvalue less than 0.05 were considered significantly enriched by differential expressed genes. The DO (Disease Ontology) database describes the function of human genes and diseases. DO pathways with corrected Pvalue less than 0.05 were considered significantly enriched by differential expressed genes. The DisGeNET database integrates human disease-related genes. DisGeNET pathways with corrected Pvalue less than 0.05 were considered significantly enriched by differential expressed genes. We used clusterProfiler software to test the statistical enrichment of differentially expressed genes in the Reactome pathway, the DO pathway, and the DisGeNET pathway.

#### Gene Set Enrichment Analysis

Gene Set Enrichment Analysis (GSEA) is a computational approach to determine if a pre-defined Gene Set can show a significant consistent difference between two biological states. The genes were ranked according to the degree of differential expression in the two samples, and then the predefined Gene Set were tested to see if they were enriched at the top or bottom of the list. Gene set enrichment analysis can include subtle expression changes. We use the local version of the GSEA analysis tool (http://www.broadinstitute.org/gsea/index.jsp), GO, KEGG, Reactome, DO and DisGeNET data sets were used for GSEA independently.

#### PPI analysis of differentially expressed genes

PPI analysis of differentially expressed genes was based on the STRING database, which known and predicted Protein-Protein Interactions.

